# Temporal-Coherence Induces Binding of Responses to Sound Sequences in Ferret Auditory Cortex

**DOI:** 10.1101/2024.05.21.595170

**Authors:** Kai Lu, Kelsey Dutta, Ali Mohammed, Mounya Elhilali, Shihab Shamma

**Affiliations:** Emory University Medical School; Electrical and Computer Engineering Department & Institute for Systems Research, University of Maryland College Park; Electrical and Computer Engineering, The Johns Hopkins University; Départment d’étude Cognitives, école normale supérieure, PSL

**Keywords:** Temporal coherence, Binding, stream formation, attention

## Abstract

Binding the attributes of a sensory source is necessary to perceive it as a unified entity, one that can be attended to and extracted from its surrounding scene. In auditory perception, this is the essence of the cocktail party problem in which a listener segregates one speaker from a mixture of voices, or a musical stream from simultaneous others. It is postulated that coherence of the temporal modulations of a source’s features is necessary to bind them. The focus of this study is on the role of temporal-coherence in binding and segregation, and specifically as evidenced by the neural correlates of rapid plasticity that enhance cortical responses among synchronized neurons, while suppressing them among asynchronized ones. In a first experiment, we find that attention to a sound sequence rapidly binds it to other *coherent* sequences while suppressing nearby *incoherent* sequences, thus enhancing the contrast between the two groups. In a second experiment, a sequence of synchronized multi-tone complexes, embedded in a cloud of randomly dispersed background of desynchronized tones, perceptually and neurally pops-out after a fraction of a second highlighting the binding among its coherent tones against the incoherent background. These findings demonstrate the role of temporal-coherence in binding and segregation.

## INTRODUCTION

Humans and other animals often perceive and manage auditory signals emanating from many simultaneously active sources in cluttered environments. The acoustic signals arrive to the ears as mixtures to be segregated, tracked, and recognized. This feat is achieved via multiple complex cognitive functions and neural processes including attention, memory, and rapid plasticity [1,2]. But a key role is played by a simpler process referred to as the *principle of temporal coherence* [1,3]. It postulates that a single source primarily evokes persistently-synchronized (or coherent) responses representing its various attributes (e.g., pitch, location, timbre). Furthermore, activations due to other independent sources are typically mutually incoherent (e.g., two simultaneous speakers uttering different words). Consequently, binding the coherent attributes of a target source would segregate (extract) it from other incoherent sources. Numerous psychoacoustic tests, EEG/MEG and single-unit auditory cortical responses [3–6] have over the last decade confirmed many aspects and surprising predictions of this hypothesis in scene analysis. A review of these studies with detailed commentary can be found in [1].

The experiments described here build upon past investigations of the neural basis of perception of simple tone sequences as “streams”, or stream formation [2]. For example, one specific study focused on auditory cortical responses as animals attended to synchronized *vs* alternating tone-sequences [7], as depicted in **Figure 1A**. The rationale is that the evoked responses mimic those due to more complex source mixtures, with neurons tuned to these tones becoming coherently (top panel) or incoherently driven (lower panel). Such activations are postulated to induce rapid plasticity of the mutual connectivity among the responsive neurons, with coherent neurons forming mutually excitatory connections (depicted in red) that enhance both of their responses and gradually bind them together. The opposite is hypothesized to occur when neurons are driven incoherently (bottom panel), where mutually inhibitory connectivity forms suppressing both neurons’ responses. The results in [7] confirmed these postulates by demonstrating that during attentive listening, responses rapidly evolved (enhanced and suppressed) with dynamics of the order of 100’s ms.

**Figure 1:**
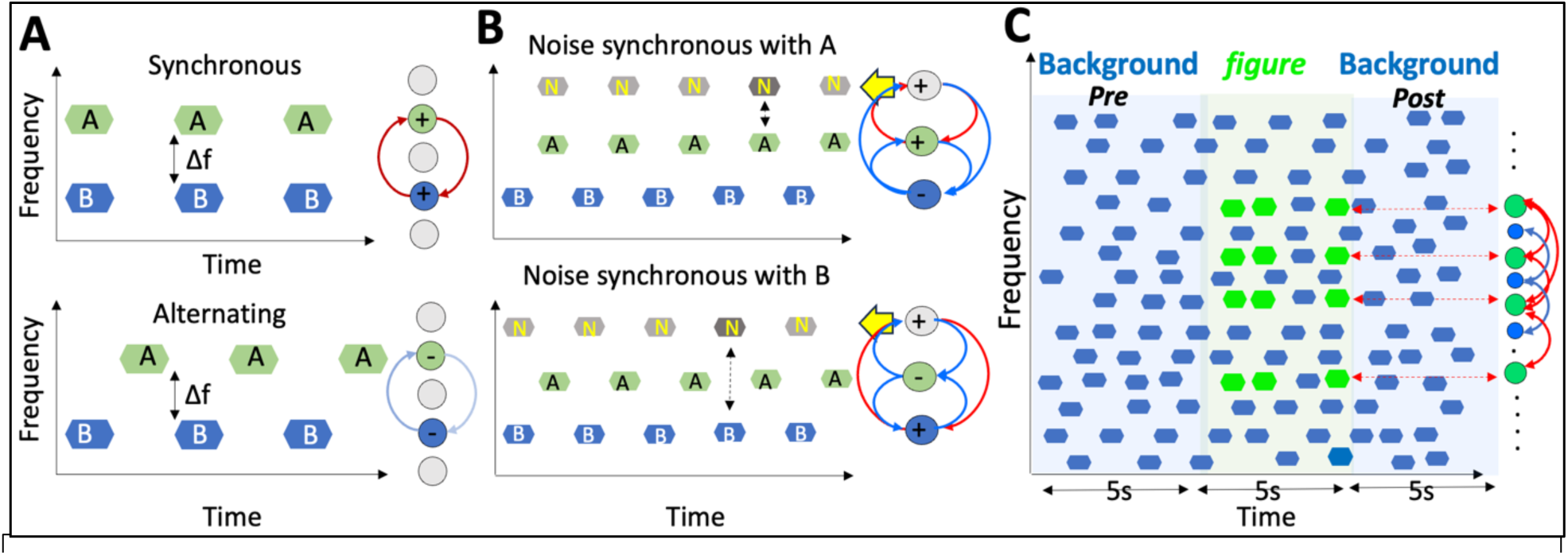
Temporal coherence and the binding hypothesis. (A) Binding in the classical two-tone streaming paradigm. Synchronous tones (**A, B** in top panel) form a single perceptual stream and induce coherently driven neurons to form mutually excitatory connections (red arrows) and enhanced responses. When asynchronous (bottom panel), the tones segregate into two streams and induce mutually inhibitory connections (blue arrows) that suppress the responses. **(B) Binding through selective attention.** Attending selectively to a sequence of noise-bursts (arrows) makes it serve as an anchor that binds it to other synchronous sequences, while asynchronous sequences become suppressed. The role of sequences **A** and **B** can be interchanged to monitor the effects on the responses of the cells tuned to the two tones. **(C) The stochastic figure-background stimulus** consists of a random tone-cloud (blue tones) with an intermediate epoch (***figure***) during which a sequence of several coherent tones (4, 6, 8, or 10) are introduced interspersed irregularly with the random tones at a rate of 4/s. It is postulated that the coherent *figure* tones become bound (red arrows), their responses enhanced, and then pop-out perceptually.

Experiment I in this study adds a fundamental twist with an auditory scene that is created with *multiple* sequences of different tokens (e.g., tones and noise) that could be perceptually organized in different ways depending on how attention is selectively deployed. **Figure 1B** depicts such a scenario where two alternating tone sequences (**A, B**) are presented with a 3^rd^ sequence of high frequency noise-bursts (**N**) which can be synchronous with either of the tone seq4uences (top and bottom panels of **Figure 1B**). The rationale is that when a listener selectively attends to the noise-burst sequence, the percept of the whole scene evolves to bind the 2 coherent sequences, i.e., **N**-**A** (*top panel*) or **N**-**B** (*bottom panel*), and to stream them apart from the remaining alternating tone sequence (see [2], p.29). How might this occur physiologically is depicted schematically by the pattern of dynamic connectivity postulated to form following the same process discussed in Fig. 1A. To test this hypothesis, Experiment I here assessed the evolution of *binding* as evidenced by the changing responses to the tone-sequences when the animal attends in one condition *versus* another (*top vs bottom panels*). It is specifically predicted that (**1**) in the top-panel condition, A-tone responses would *increase* while **B**-tone responses *decrease*; the opposite would occur in the bottom panel. Another prediction is that (**2**) evolution of the connectivity (and response changes) is directed or controlled by the attentional focus in favor of the tone sequence synchronous with the noise. Consequently, in the absence of attention (passive listening), this binding process and its concomitant suppression or enhancement are predicted to weaken or disappear.

In Experiment II, the number of coherent tones and the complexity of the overall auditory scene is increased significantly (Figure 1C), while remaining relatively accessible and interpretable within the same hypothetical scenarios of Figures 1A**, 1B**. Here a random cloud of incoherent tones (referred to as the “Background”) is heard typically for 5 seconds, followed by an additional embedded sequence of coherent tone-complexes (referred to as the *figure*) that commences abruptly, and then repeats *irregularly at* about 4 per second for a total of 5 seconds. The Background tone-cloud continues for a final 5 seconds (post-*figure*) interval (Fig. 1C). It is hypothesized that the coherent tones of the *figure* will induce excitatory connectivity among the coherently driven neurons, binding them together and enhancing their responses, and thus making the *figure* perceptually pop-out, becoming more salient after a short period of buildup. This type of stimulus (often referred to as the *Stochastic Figure Ground (SFG)*) has already been fruitfully investigated with human subjects in numerous psychoacoustic and MEG/EEG studies [8–10]. But no single-unit recordings of the underlying responses in an animal model have been reported.

The results reported here from the two Experiments of Figs. 1B, 1C are broadly consistent with the idea of a binding process in which coherent neural responses become rapidly enhanced relative to the suppression of incoherent responses. In **Experiment I**, the effect of selective attention reshapes the organization of the responses to the two-tone sequences (Fig. 1B). In **Experiment II**, responses in a passively listening animal exhibit the consequences of binding on the coherent tones of the *figure* (Fig. 1C), in agreement with findings from EEG human recordings passively listening to similar kinds of stimuli [10]. Finally, we employ **f**unctional **U**ltra**S**ound (fUS) imaging of the cortical responses to view large-scale spatiotemporal dynamics of brain activity during coherent activation and thus gain a multi-scale perspective of the binding process.

## RESULTS

A total of 4 ferrets participated in all experiments. Two ferrets (**U,R**) were involved in **Experiment I**, and three (ferrets **B,K,U**) in **Experiment II**. Single-unit recordings were made in the primary auditory cortex (A1) unless stated otherwise. *fUS* recordings spanned a cortical region encompassing the medial and posterior ectosylvian gyri in ferrets **K,U** (primary, anterior, and secondary (PEG) auditory cortical fields).

### Experiment I: Binding and selective attention

Ferrets **R** and **U** listened to simultaneous streams of sound sequences (Fig. 1B) consisting of a high-frequency band of noise-bursts, and two lower frequency tone sequences (**A** & **B).** The animals were trained to attend *only* to the noise bursts (**N**) and to detect a small intensity change (denoted as a darker burst in Fig. 1B) to receive a water-reward. The center frequency of the noise burst was selected within a 5-octave range of the tones. Unbeknownst to the animals, two tone conditions were tested where either of the **A** or **B-**sequences were synchronous with the noise bursts (*upper* & *lower panels* of Fig. 1B). Animals generally performed the task identically in the two conditions as exemplified by the matched performance of ferret **R** during the two sets of trials (Supplementary Figure 1S). Single-unit responses were recorded in ferrets **R** (76 neurons) and **U** (40 neurons) in both *passive* and *active* conditions so as to compare the effects of engagement and attention. In each recording, we first measured the best frequencies (BF’s) of all simultaneously isolated neurons (on 4 independent electrodes). Then the frequency of one of the two tone-sequences was selected near the BF of one neuron, while the frequency of the other tone was selected 3/4 octave away and nearer to one of the other isolated neurons. This way, at least some neurons preferentially responded to either tone **A** or **B** in each recording.

Two types of trials were used to stimulate each neuron (e.g., the green neuron in Fig. 1B whose BF is aligned with **tone A** sequence): (1) *Coherent (or Synchronized - **SYN**) trials* in which the neuron is driven by **tone-A** synchronously with the noise stream (N); (2) *Incoherent (or Asynchronized - **ASYN**) trials* in which the BF-tone of a cell (**tone A**) is asynchronous with the noise, while the other tone (**Tone B**) drives the blue neuron synchronously with the noise bursts. We recorded responses of *each* neuron to *both* stimulus conditions while the animals passively listened and when they were actively engaged in the detection task. The neural responses from each pair of (**A** and **B**) tones are *averaged* over all their repetitions in each condition, and also over the duration of all trials. The two PSTH’s thus generated from each neuron are labeled according to the trial type as: ***SYN_pass_*** *, **SYN******_act_****, **ASYN******_pass_*** *, **ASYN******_act_***. A trial typically lasted a minimum of 1.5s (4-20 repetitions of the **A**, **B** tone pairs) to allow for the postulated neuronal connectivity to adapt during the trial and become evidenced by the changes (enhancements or suppression) in the single-unit responses. Note that since a trial-type was chosen randomly, the changes observed are assumed to buildup from scratch at every trial.

#### Evidence of binding in the patterns of rapid plasticity

Neuronal responses from a given cell depended on many factors, and hence were quite variable. They included the exact frequency of the **A** and **B** tones relative to the BF, and the excitatory and inhibitory fields of the cell’s Spectrotemporal receptive fields (STRFs). Another important factor is the behavioral state of the animal (passive or active) because it has been commonly found that when animals engage in a task and attend to its stimuli, the overall responsiveness of primary auditory cortical cells often decreases significantly [22,23,35]. Therefore, to demonstrate the binding hypothesis based on such responses, it is critical to consider the *relative* changes of a cell’s responses between the SYN and ASYN conditions and not simply whether it is enhanced or suppressed in absolute terms as we illustrate next.

Figure 2A panels displays PSTH’s from two neurons in ferret **R.** Each panel depicts the average PSTH responses in the SYN (left-half = 320 ms) and ASYN (right-half = 320 ms) conditions as illustrated by the tone and noise burst symbols above the panels. The two cells’ PSTHs are quite different yet both are consistent with the binding hypothesis. *Unit-1* (*top panel;* Fig. 2A) exhibits responses that follow closely the postulate described in Fig. 1B – with attention during the task, SYN responses (left-side) increase while the ASYN responses decrease relative to the passive state. We quantify these changes as **D** and **Δ** in ***Equation 1***:

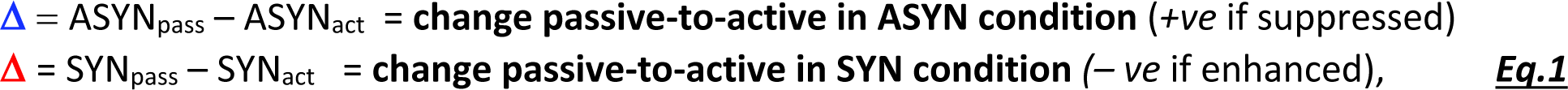

where *SYN_pass_, SYN_act_, ASYN_pass_, ASYN_act_* denote the *mean of PSTH* responses over the 320 ms interval comprising the two tones depicted in each panels of Fig. 2A. Note that according to **Eq.1**, **D > Δ** in this neuron because ASYN_pass_ > ASYN_act_ (**D** > 0) and SYN_act_ > SYN_pass_ (**Δ < 0**).

**Figure 2.**
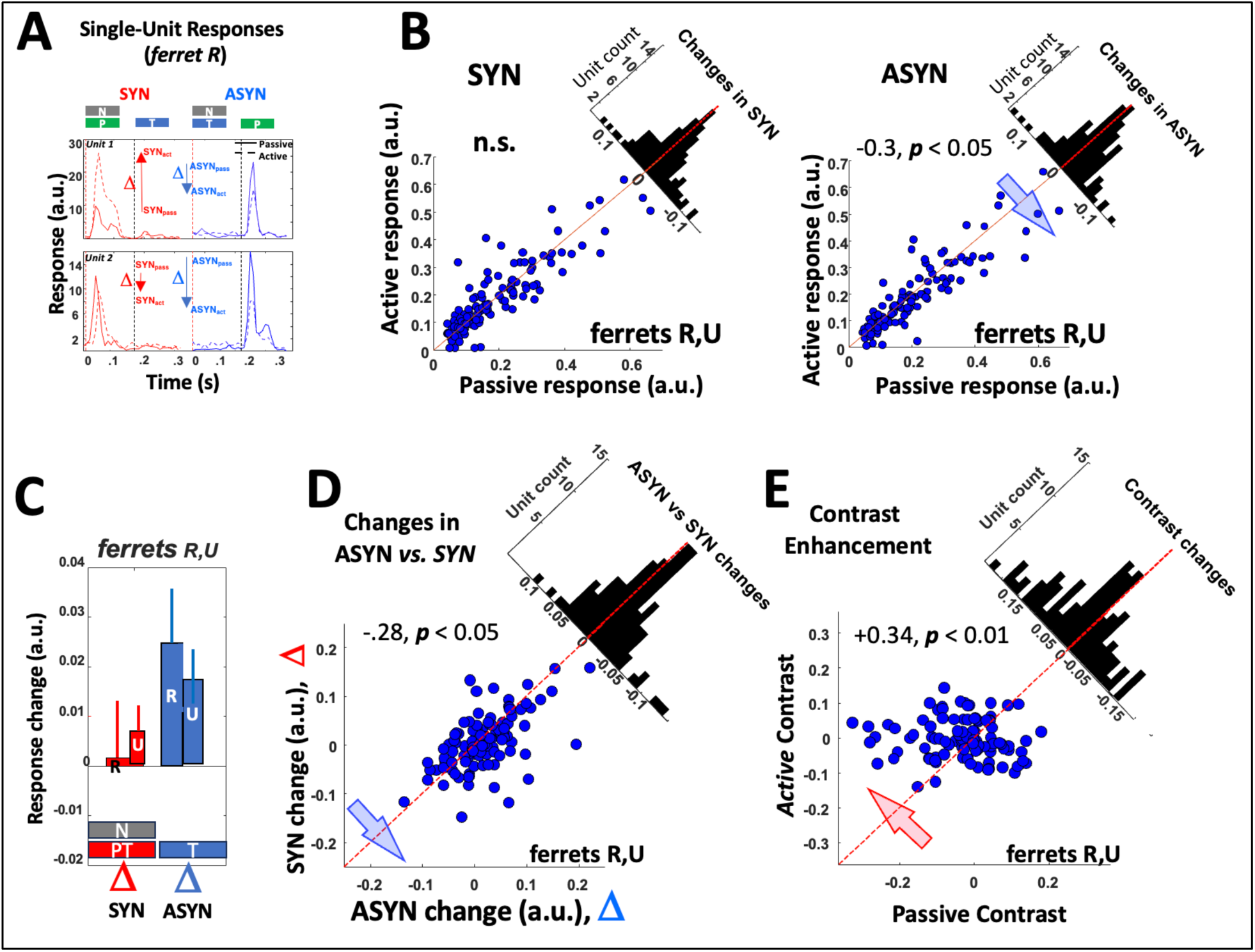
Response modulation reflects binding of different tone streams. **(A) Enhancement and suppression in single cell examples in ferret R**. In each of the (*top and bottom*) panels, the average PSTH of the SYN (*left-half*) and ASYN (*right-half*) trials. The stimuli are indicated by the colored symbols on top. The noise-burst (N) sequence is synchronous with either of the two tone-sequences - the preferred tone (P) usually chosen near BF, or with the non-preferred tone T. *Unit-1* PSTH responses (*top panel*) change during the active state (dashed-curves) relative to the passive state (solid-curves). the *inset* schematic displays the level of the averaged responses in the four different conditions (SYN_act_, SYN_pass_, ASYN_act_, ASYN_pass_). This cell’s responses are enhanced (red arrow, **Δ**) when driven synchronously with N, and suppressed when asynchronous (blue arrow, **D**). Both **Δ** and **D** are defined as the *<u>change</u>* <u>from *passive to active*</u> *(Eq.1 in text).* Therefore, for *unit-1* Δ < 0 and **D > 0**, or **D** > **Δ**. In *unit-2* (*bottom panel*) both SYN and ASYN responses are suppressed in the active state, but the suppression is stronger in ASYN, and hence **D** > **Δ**. **(B) Scatterplots of response modulations in Syn and ASYN conditions in both ferrets R & U**. (*Left panel*) The scatterplot of the average responses of each unit in SYN trials during the active condition (Y-axis) versus passive (X-axis). (*Right panel*) The scatterplot of the same cells during the ASYN trials. The histograms summarize the scatter around the midline, and its effect-size is indicated if significant (see test & **Methods**). Only during the ASYN trials is there a significant bias of responses *below* the midline (suppression). **(C) Suppression is strongest during the ASYN trials** in both ferrets, as indicated by the bar-plots of the means and standard errors of spike rate differences (*Pass – Act =* D, **Δ**) in both trial types. **(D) Evidence of binding in scatterplots of response modulations from all cells in ferrets R** & **U.** Scatterplot of response changes during ASYN (**D**) versus SYN (**Δ**) indicates a significant bias below the midline (effect size = -0.28, *p* < .05) implying an overall **D** > **Δ**, taken to be as evidence of binding. (E) Contrast enhancement due to binding. *Normalized* differences between SYN and ASYN (or the contrast) increases during the active state, as indicated by the significant bias of points above the midline (effect-size = +0.34, *p* < .01).

Because of the depressive effects of task engagement and other factors alluded to above, it is common in cells’ responses for the inequality **D > Δ** to hold even when both conditions cause a suppression of the responses as in *unit-2* (*lower panel*; Fig. 2A). This cell showed deeper suppression during ASYN compared to SYN, hence **D > Δ**, which is still in line with the binding hypothesis.

Figure 2B summarizes the response changes over all neurons in both ferrets (**U**,**R**). The two panels contrast the response modulations between passive *vs.* active states, separately under SYN (*left panel*) and ASYN (*right panel*) conditions. A significant suppressive change occurs in the ASYN condition, as indicated by the negative *effect-size* and ***p***-value (see **Methods**) as the animal attends to the noise stream (Fig. 1B). In the SYN condition, many cells are no longer suppressed, but there is no significant positive bias either. This suggests that the binding effects are rooted more in the *suppression* of unattended ASYN responses, or in the *relative* change between responses in the passive and active conditions. The net response changes in all cells from both ferrets (**R,U**) are integrated and depicted by the two bar plots in Figure 2C which show a strong net suppression during ASYN and a weak net increase in SYN. All above findings were replicated individually in each of ferrets **U** and **R** (***Supplementary Figures 2SA-2SB***).

Figure 2D directly compares the response changes (**D** *vs.* **Δ**) in each cell due to engagement. The negative bias confirms that the inequality **D > Δ** predominates, providing evidence of binding induced changes. However, despite the changes in SYN (**Δ**‘s) are relatively small and unbiased (Fig. 2B**, 2C**), they apparently still contribute to the overall plasticity patterns reflecting binding. To demonstrate this, we shuffled the list of **Δ**’s associated with all cells, hence scrambling the pairing of **D** with **Δ** in each cell, and then recomputed the scatterplots of Fig. 2D. The **Δ** shuffling disrupts the condition **D > Δ** in many cells and eliminates the earlier negative bias (***Supplementary Fig. 2SC***).

#### Evidence of binding through contrast enhancement

Another important consequence of binding’s competitive and cooperative interactions (Fig. 1B) is to *increase the contrast* between the responses of the attended coherent cells *relative* to the incoherent ones. To test if this change occurs in our data, we defined for each cell the average contrast between the SYN and ASYN responses in passive and active conditions as:

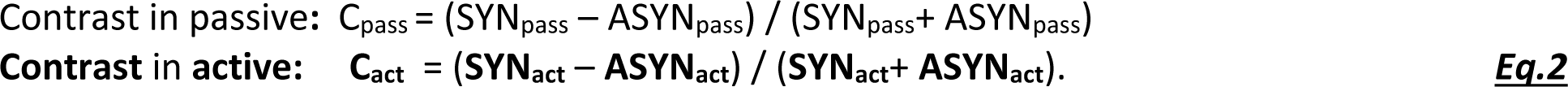

These normalized contrast measures are compared in the scatterplot of Figure 2E, which displays a significant *positive* bias with **C_act_** > C_pass_ (+0.34, ***p***<.01) implying that task engagement increases the contrast between the coherent *relative to* the incoherent responses. Finally, to demonstrate the importance of **Δ**‘s to contrast enhancement, we find that removing their contributions by setting **Δ**=0 (SYN_act_=SYN_pass_) abolishes the positive shift of Fig. 2E, as demonstrated in ***Supplementary Figure 2SD***

In summary, the results thus far reveal that selective attention to a sequence induces response changes consistent with the binding hypothesis (**D > Δ**), but which take the form of larger suppression **D** relative to **Δ** effects. Nevertheless, manipulating (shuffling or nulling) the **D**‘s has profound effects on the change distributions and contrast indices reflecting their importance.

### Experiment II: The stochastic *figure*-ground stimulus

This experiment employs the stochastic *figure*-ground (SFG) stimulus (Fig. 1C), a versatile stimulus that allows for multiple simultaneous measurements including the neurons’ STRFs, and the dynamics of their response changes as the *figure* exerts its strong influence by inducing binding among the coherently driven neurons. The stimulus consists of three epochs: **(i) Pre:** the *initial* epoch of random tones that facilitates STRF estimation (referred to as *pre-*STRF); **(ii) Mid**: an *intermediate* period that contains in addition to the random tones several irregularly repeated bursts of temporally coherent tones – the *figure* – at about 5 bursts per second. It is postulated that during this epoch binding evolves among the coherently driven neurons; finally **(iii) Post:** another epoch of random tones that allows for a *post*-STRF measurement and assessment of the persistent effects of the sustained presentation of the synchronized *figure* tones during the intermediate epoch.

Note that the timing of the random and coherent tones throughout this stimulus was chosen carefully to ensure that when added together, the overall stimulus envelope did not exhibit large fluctuations related to the synchronous tones of the *figure*. Specifically, as elaborated upon in **Methods**, the tones in the random tone epochs were overlapping and had roughly the same overall power throughout the *pre-* and *post-figure* epochs. Importantly, during the *figure*, the coherent tones were created by adjusting the timings of nearby random tones to keep the overall number of tones at any given moment roughly constant. Furthermore, we also ensured that the synchronous onsets of the *figure* tones were balanced by removing any random tones that have nearby onsets. Three ferrets participated in the single-unit recordings of this Experiment. Details of the numbers of isolated single-units, shared ferret participation across different Experiments, and their ages are available in **Methods**.

#### STRF measurements during the random tone epochs

The random tone stimulus is akin to a noise that can be used to measure the STRFs and hence the tuning and expected dynamics of the cells [12–13,19]. We first validated that this stimulus could recreate meaningful STRFs that resemble those due to our standard battery of measurements in A1, specifically using *temporally orthogonal ripple combinations* (TORC) stimuli [13]. With lag values of -10 to 120 ms, the cross-covariance between the response and the auditory spectrogram of the stimulus was computed and normalized by the auto-covariance of the stimulus. Ridge regression was performed with 30 log-spaced values for each trial and then averaged to compute the *pre-*STRF and *post*-STRF for an interval of 1 or 2-second just before and after the *figure* epoch. During the *figure* intermediate epoch, we computed three additional similarly windowed STRFs during the *early*, *mid*, and *late* portions of this epoch. While computed the same way as the *pre-*STRFs and *post-*STRFs, these measurements are different in that the stimulus auto-correlation is not white when the *figure* is included. The resulting artifacts due to the *figure* are quite evident, but the ‘STRF’ estimates nevertheless are revealing and useful as we explain below.

Figure 3A compares the STRFs measured with the random tone *versus* TORC responses (ferret **B**). The two differ since many factors influence such estimate [12,13] including that TORCs contain a dense range of frequencies spanning 5 octaves which evolve over time in a continuous manner. The random tones by contrast contain only a discrete set of 37 frequencies and are presented as discrete tone sequences spanning 3 (or often 6) octaves, and hence the striated appearance of the excitatory regions of the bottom-rightmost panel of Figure 3A. Despite these large differences between stimuli, the two measurements display approximately matched latencies, BFs, and shapes of the major excitatory or inhibitory regions near the BF. The random tone STRFs are particularly useful in monitoring how binding evolves over the different epochs, and revealing the effects of the *figure* presentation during the intermediate epoch.

**Figure 3.**
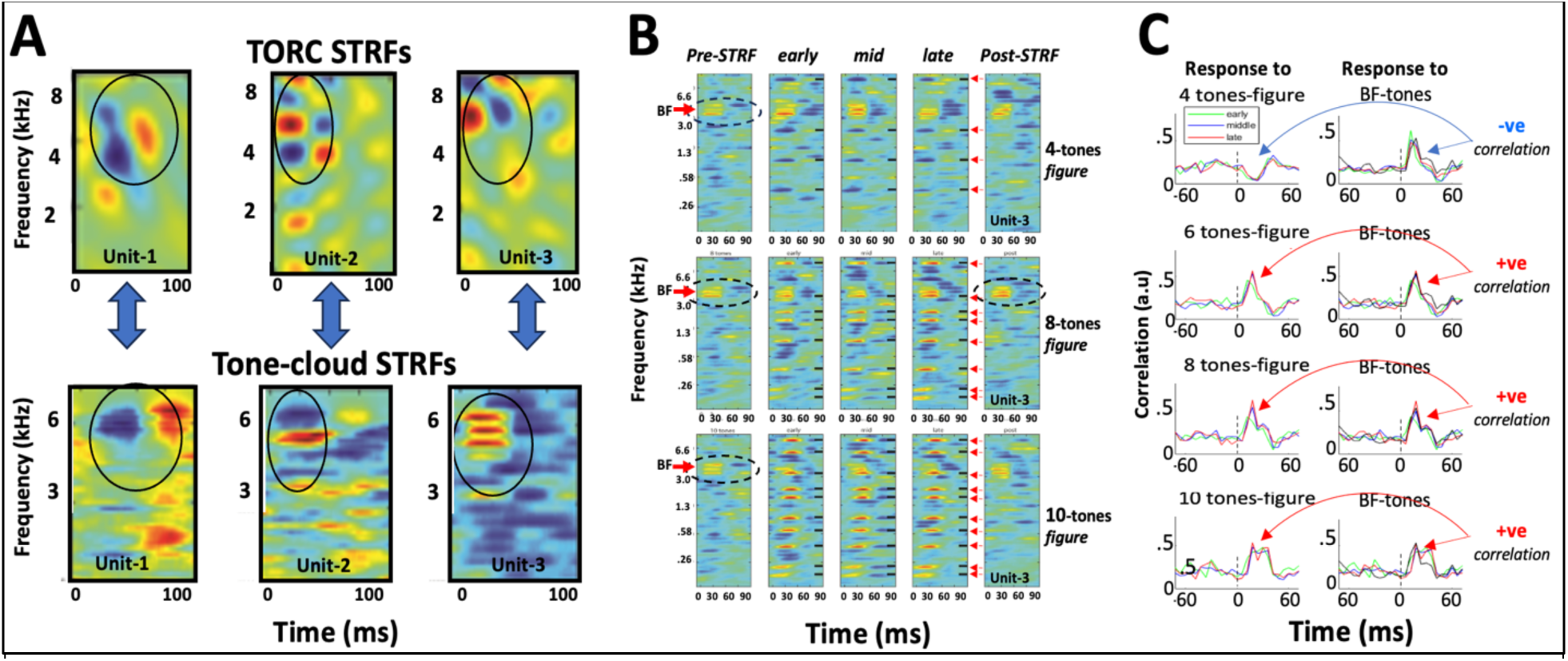
STRF measurements in relation to the *figure* in the SFG. (**A**) **Comparing STRFs measured with TORCs and random tones.** Examples of 3 units where TORC STRFs (top row) share prominent features with tone-cloud STRFs (bottom row) but differ in detail. **(B) STRFs of a single cell measured in different epochs of the stimulus.** *Pre*- and *post-*STRF are computed from the pure random tones, with the dashed circles highlighting the BF of the cell. In the intermediate epochs (*early*, *mid*, and *late*), the STRFs exhibit the effects of the *figure* presentation through the negative (blue) and positive (red) striations (see text for details). **(C) Measuring the correlation between the *figure-* triggered and BF-triggered responses.** Onset of *figure* and BF tones-triggered average responses are at time=0 ms. When a *figure* drives a cell (*left column plots*) in a similar way to the BF tones (*right column plots*), the two PSTHs are positively (***+ ve***) correlated (as in the 6-, 8-, and 10-tone *figures*). Negative correlations (***-ve***) are seen for the 4-tone *figure* (*top row of plots*).

### STRFs during *figure* presentations

Figure 3B illustrates a series of 5 STRFs computed from the responses to *unit-2* (Fig. 3A): a *pre*-*STRF*, 3 *intermediate-epoch (early-, mid-, late-STRF),* and a *post*-STRF. Up to four different *figures* were presented in a randomly interleaved manner: 4 tones (*top row*), 6 tones (*not shown*), 8 tones (*middle row*), and 10 tones (*bottom row*). The frequencies of the *figure* tones used in each panel are indicated by black-markers along the rightmost edge of each panel, and the red-arrows along the rightmost edge of the *late-STRF* panels. As is evident, the *pre-STRF* and *post-STRFs* resemble each other and exhibit a clear BF (highlighted by the dashed oval circles). Enlargements of some of these panels are shown for added clarity in **Supplementary Figure 3S**.

STRFs in the intermediate (*figure*) epoch display strikingly different features of either *suppressed* (blue stripes in the 4-tone panels) or *enhanced* (red/yellow stripes in the 8 & 10-tone *figure* panels). These narrowly-tuned features emerge because the *figure*-tones are exactly synchronized (strongly-correlated), and hence the stimulus is not spectrally white. Consequently, if one of the tones of the *figure* activates the cell near its BF at 6 kHz (e.g., the 2^nd^ tone from the top in the 8-tone *figure,* or the 3^rd^ tone in the 10-tone *figure*), then the responses will be highly correlated with all the other *figure*-tones resulting in an STRF with enhanced striped regions (*middle and bottom row panels* of Fig.3B). A valuable insight gained from the emergence of these extra peaks is the evidence that the *figure* in fact activated the cell with one of its tones. When none of the *figure*-tones activates the cell, or if they suppress the cell’s responses as when aligned with its inhibitory sidebands, the *figure*-tones become negatively correlated with the cell’s responses, and thus induce suppressed (blue striped) regions in the STRF (*top panels* of Fig.3B). Again, one can infer here that unlike with the 8- and 10-tone *figures*, the cell was incoherently activated (asynchronous) relative to cells that are positively driven by the *figure*.

In the interest of avoiding additional nomenclature, and since we use precisely the same algorithms to compute the *pre*-STRF and *post*-STRFs as we do during the *figure* presentations, we shall continue to refer to these measurements as STRFs (*early*-, *mid*-, *late*-) although we are aware of the limitations of these estimates as strictly STRFs. Furthermore, because of the noisy character of all the STRFs, the estimates are smoothed by removing all but the most significant voxels (as detailed in **Methods**) and then summing the remaining significant voxels to give an estimate of the ‘strength’ of the STRF, or indirectly the cell’s responses. Since the *pre*-STRFs and *post*-STRFs are measured with the same random tones, comparing them allows us to directly estimate the STRF changes due to the intervening *figure*. By contrast, *early*-STRFs, *mid*-STRFs, and *late*-STRFs reveal the *dynamics* of STRF changes during *figure* presentation, and how the *figure* activations of a cell relate to its overall effects on the STRFs.

The effects of the *figure* on cortical cells are exemplified by the responses and STRF changes in the unit of Figure 3B. For instance, the detailed features of the *early*-STRFs, *mid*-STRFs, and *late*-

STRFs are dependent on the arbitrary experimental alignment of the *figure* relative to the BF. Thus, on the one hand, the cell is *suppressed* by the 4-tone *figure* (*top panels*; Fig.3B) inducing inhibitory responses aligned to the *figure* tones. Consequently, this cell’s STRFs exhibit gradually diminished BF peaks (*early **>** mid **>** late* STRFs) and an overall no-change between the *pre*- and *post*-STRFs (to be quantified later). On the other hand, the same cell is positively-driven by the 8- and 10-tone *figures* (*middle and bottom rows of panels*), causing coherent responses that seem to enhance its STRFs. This condition is likely to occur with bigger *figures* as they are more likely to contain tones that align with the BF’s.

To capture and summarize STRF changes from a large population of cortical cells, it is essential to consider the alignment of the *figure* tones with the cells’ BFs for *each cell* and *test* separately. Figure 3C illustrates how we quantify this relationship by first computing the ***figure*-triggered PSTH** as illustrated by the *left column of panels*. These clearly indicate triggered-PSTHs that are suppressed for the 4-tone *figure*, and excitatory for the 6, 8, and 10-tone *figures*. We then measure the BF-triggered PSTH of the cell during the *pre-* or *post-figure epochs* (*right column of panels*). The match (inner-product) between the two PSTHs is indicative of the nature of the *figure* activations: if the match-sign is negative (***-ve***) as in the 4-tone *figure* case (*top row*), it indicates the presence of blue-stripes in the *early-*, *mid*-, and *late*-STRFs of the corresponding panels in Fig. 3B; if the sign is positive (***+ve***), it reflects the presence of red-stripes during the *figure*.

#### STRF response changes and binding in the neuron population

We consider next STRF changes due to *figure* presentations in a large population of diverse cortical cells. When a *figure* aligns with the BF in a portion of cortical cells, it stimulates coherently these cells and induces **+ve** correlations (Fig. 3C). The proportion of such cells becomes larger with bigger *figures* because there are more tones that may align with the cells’ BFs. Since coherent responses are hypothesized to bind the responsive cells (Fig. 1B), we expect to detect more enhanced responses with bigger *figures,* which in turn leads to the perceptual “pop-out” of the *figure* [14].

To test these hypotheses, we first examined response changes as a function of *figure* size in a large population of cells. Figure 4A displays the distributions of the STRF changes between the *pre*- and *post*-STRF in recordings from *all* 3 ferrets (1668 tests in 277 cells), clustered according to the size of the *figures*. For each test, the STRF change is defined as:

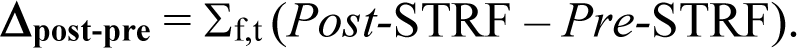

As predicted earlier, the distribution of the 10-tone *figure* changes is significantly more positively shifted *(**p**<.001)* relative to the 4- and 6-tone *figures* whose *means* are often insignificantly shifted.

**Figure 4.**
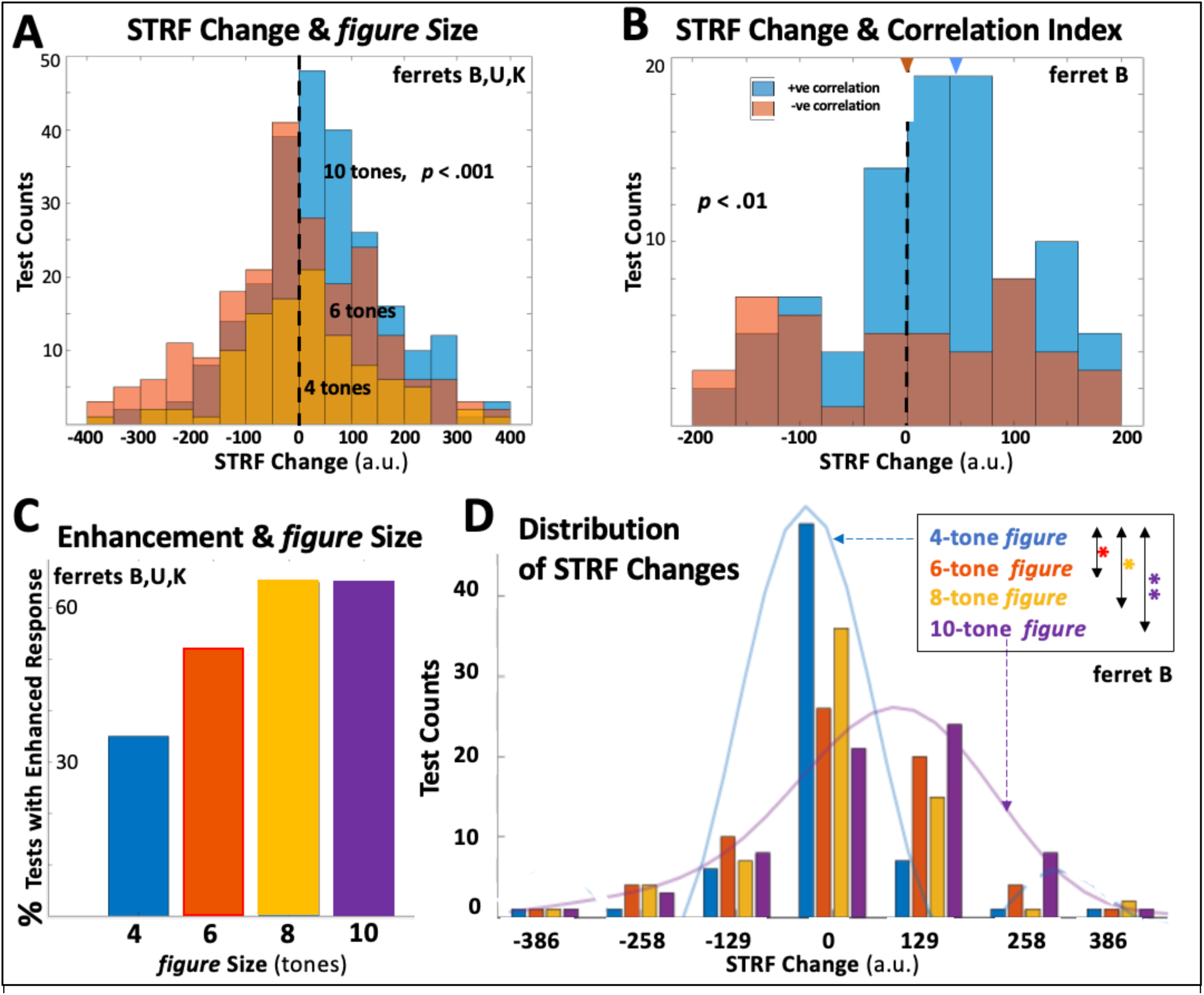
The *figure* size and the distributions of STRF changes. **(A) Distribution of all STRF changes.** All cells are included in this distribution regardless of their correlations with the *figures*. STRFs are more likely to be enhanced with bigger *figures* (10-tones), and least for the smallest *figure* (4-tones). **(B) Distributions of STRF changes for tests with only 8- and 10-tone *figures*, clustered according to their correlations.** STRFs from +ve correlated tests are typically enhanced. Tests from - ve correlated tests are randomly distributed. **(C) Proportion of tests with enhanced responses** as a function *figure* size. **(D) Distribution of all STRF changes** as a function of *figure* size. STRF changes for larger *figures* are more significantly positively shifted.

To confirm that the alignment of the *figures* with the BF can make a significant difference to the STRF modulations, we combined in Figure 4B the tests of 8- and 10-tone *figures* in ferret **B** (125 tests) and clustered them according to their positive or negative correlation signs. As postulated, cells with **+ve** correlations exhibited significant STRF enhancements (blue distribution) compared to the centered distribution of the **-ve** correlated cells. Therefore, when cells respond coherently to a *figure,* their responses and STRFs become more enhanced, hypothetically because of the cooperative binding among them (Fig. 1C).

A more succinct view of STRF *enhancements* in all cells and tests is shown in Figure 4C where bar heights indicate the increasing proportion of enhanced cells with the *figure* size. It is also evident that 8-tone *figures* are on average as effective as the 10-tone *figures.* Finally, the full range of STRF changes with all *figure* sizes are shown in Figure 4D where all tests were conducted with the identical parameters in ferret **B** (1166 tests). Again, the 10-tone *figure* induced significantly more enhanced STRF changes (rightward shifts in the distributions; ***p****<.001*), less so for the 6- and 8-tone *figures* (***p****<.01*), and no apparent net STRF changes for the small 4-tone *figures*. Interestingly, there are no substantial *net suppressive* STRF changes (leftward shifts in the distributions) likely because of the balanced mix of **+ve** and **-ve** correlated cells even for the 4-tone *figures,* causing the overall STRF changes to cancel out. Another possible reason is that STRF changes (**Δ_post-pre_**) represent the persistent effects *after* the end of the *figure*, and it is conceivable that suppressive effects do not persist as well as enhancements. These possibilities are explored next through the dynamics of STRF changes *while figures* are presented, i.e., changes relative to the *early*-, *mid*-, and *late*-STRFs.

#### Dynamics of STRF modulations during *figure* presentations

STRF changes between the *pre-figure* & *post*-*figure* measurements (**Δ_post-pre_**) presumably buildup gradually during *figure* presentations. They can be directly captured by tracking the STRFs in each of the *early-, mid-,* and *late-* epochs. We begin by quantifying the STRF strengths as:

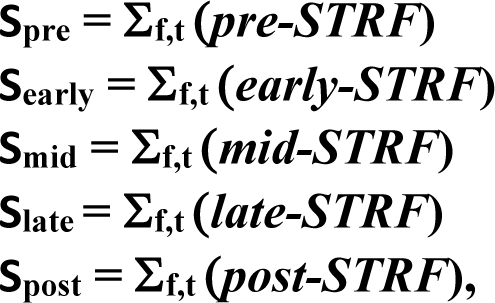

where **S** represent the integrated sum over all the *significant pixels* in the indicated STRF’s. All are computed over 1-second intervals within the different epochs (see **Methods**). The bar plots of Figure 5A illustrate the dynamics of the average STRF buildups estimated from 172 cells (371 tests) in ferret **B** and sorted according to the size of each *figure*. We note two characteristics of the responses: **(i)** during the early epoch, **S_early_** becomes significantly enhanced (relative to the initial **S_pre_**) only for the larger 10-tone *figure.* By the *mid*-epoch, within 2-3 seconds after the onset of the *figure,* the 6-, 8-, and 10-tone *figures* **S_mid_** becomes significantly enhanced. **(ii)** During the *late*-epoch, at about 4 seconds from the onset of the *figure*, the **S_late_** begins to diminish. Nevertheless, after the end of the *figure*, STRF enhancements apparently persist into the *post*-*figure* period, resulting in net positive changes between *pre*- and *post*-STRFs (**Δ_post-pre_**).

**Figure 5.**
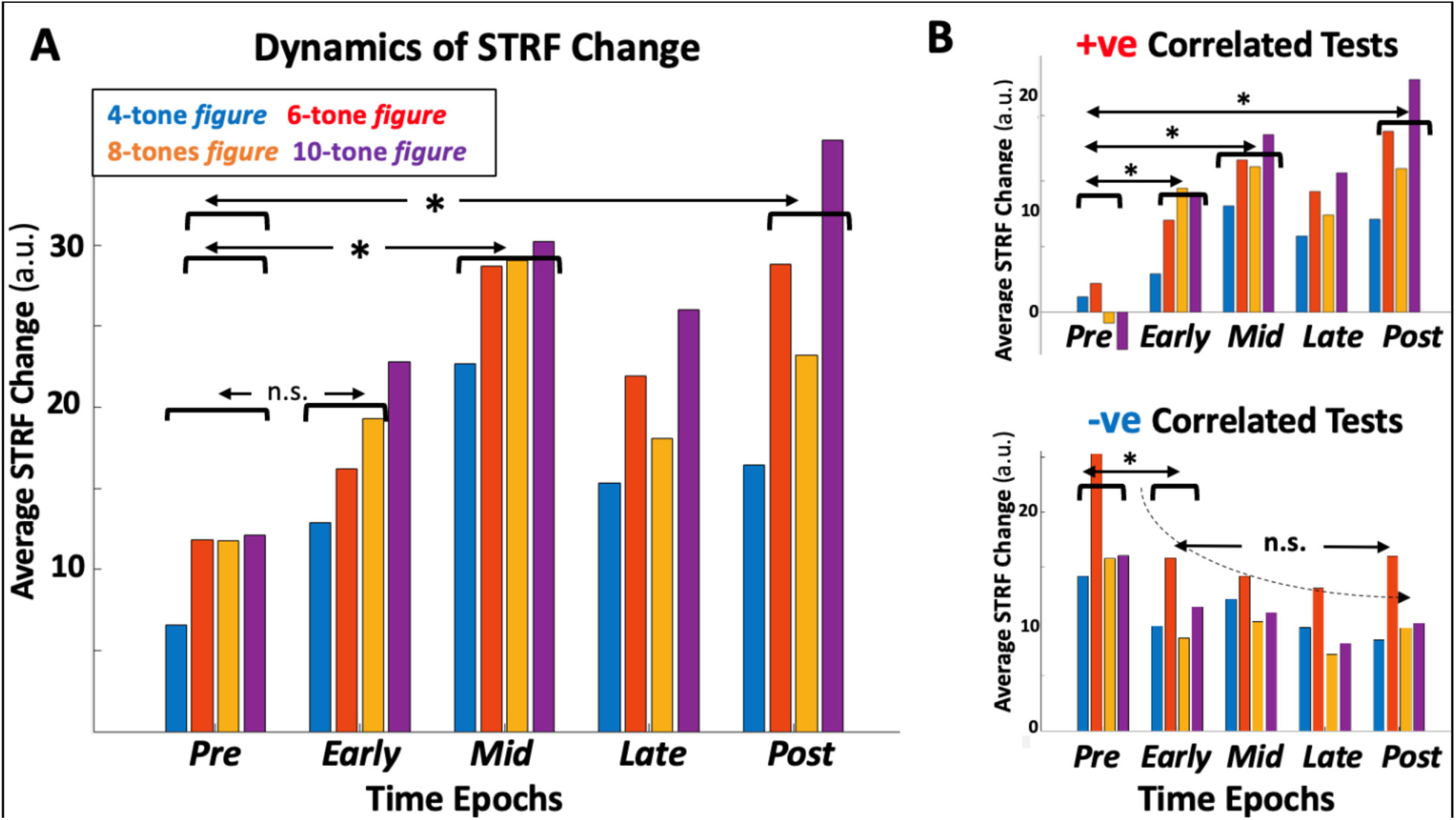
Dynamics of STRF modulations in ferret B responses. **(A) Larger *figures* modulate STRFs rapidly.** For example, the STRFs in rapidly change reaching a peak by the mid-epoch about 2 seconds from the onset of the *figure*. *Late*-STRF enhancements decrease, but they remain enhanced in the *post*-STRF relative to the *pre*-STRFs, giving evidence of the exposure to the *figures*. **(B) STRF modulations in +ve & -ve tests**. Rapid and significant STRF modulations occur, especially between the *post*- and the *pre*-STRFs during +ve tests. However, STRFs in -ve tests remain depressed throughout the tests.

STRF changes due to the *figure* activations may in fact be bigger and faster than indicated in Figure 5A because the tests included a mix of both **+ve** and **-ve** correlated tests. In Figure 5B, we segregated the two contributions. In the *top panel*, only **+ve** correlated tests are included, and all 6-, 8-, and 10-tone *figures* rapidly modulate up the cells’ STRFs during the *early*-epoch. As before (Fig. 5A), the enhancements diminish during the *late*-epoch but persist after the end of the *figure* to give a significant **Δ_post-pre_**. Interestingly, in tests with -**ve** correlations (*bottom panel* of Figure 5B), the STRFs rapidly *diminish* during all *figure* presentations, resulting in a suppressed

**Δ_post-pre_**. We conjecture that this decrease may reflect that **-ve** correlated cells are inhibited by the *figures* (e.g., 4-tone *figure* in Fig.3B), and hence they do not bind without activation. Furthermore, during such tests, cells are incoherently activated relative to the **+ve** cells, thus competing with them and becoming suppressed (hypothesis of Fig. 1C).

#### Functional ultrasound imaging of *SFG* responses

The spatiotemporal distribution of neuronal activity in the auditory cortex during an SFG trials, was tracked using **f**unctional **U**ltra**S**ound Imaging (fUS) in two ferrets (**U, Z**). This technology offers stable images of cerebral blood volume changes over large brain areas, which are assumed to reflect indirectly 1-2s delayed neuronal responses due to the random tones and *figure* over the stimulus epochs of Fig. 1C. Details of the technical and physiological procedures are available in **Methods** and previous publications [15,30–32]. Figure 6 illustrates images from ferret **U** who also provided single-unit responses in Experiments I and II.

**Figure 6.**
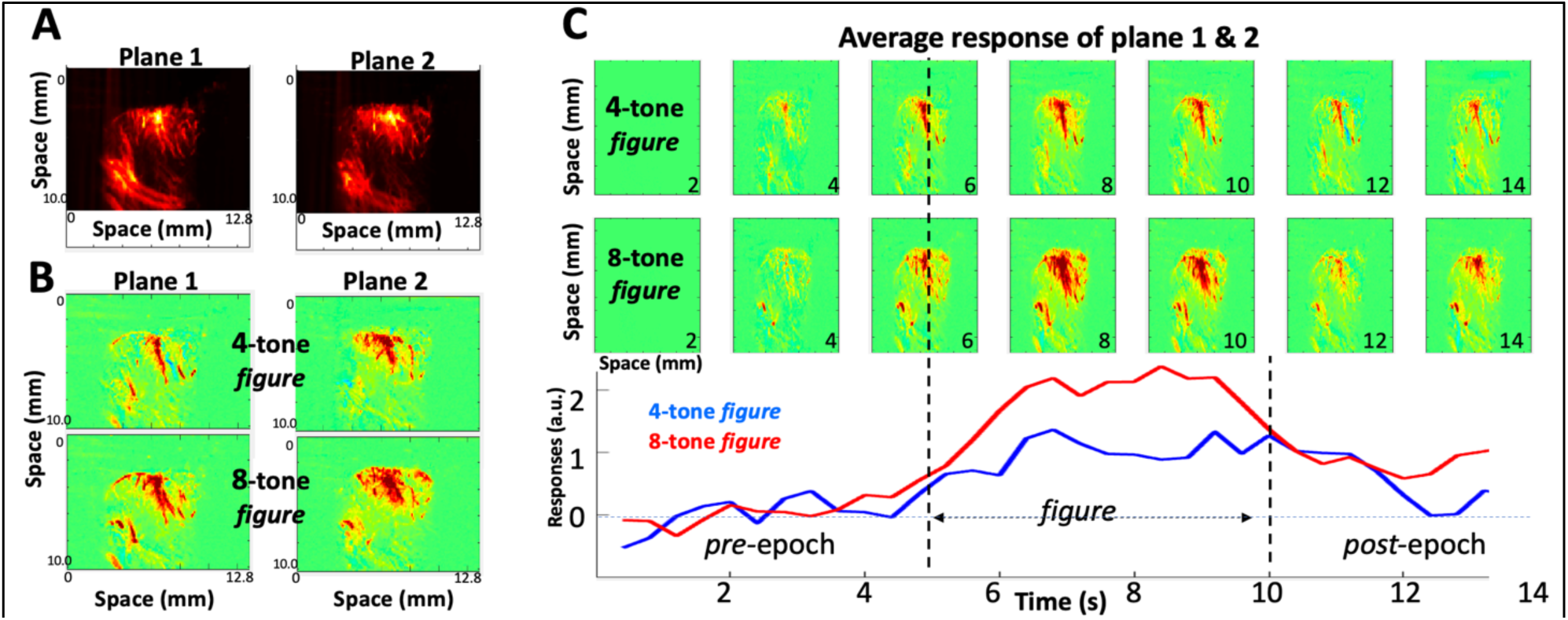
Functional Ultrasound (fUS) imaging in ferret U. (A) Anatomical vasculature of the auditory cortex. It is revealed by the activity induced during the early random tone-cloud. Two planes of the auditory cortex are imaged located at the center of the primary auditory cortex, spanning 350-500 μm in thickness. **(B) Activations averaged over the entire *figure* presentations.** The smaller 4-tone *figure* evokes significantly weaker activations than the 8-tone *figure* in both cortical planes. **(C) Dynamics of the response modulations.** Sequence of 2-sec average activations throughout the trials. Enhancements of activity are pronounced when the *figure* is large (8-tone *figure*), peaking at about 3 seconds from onset of the figure, decaying in the latter part of the *figure* and post-*figure* epoch.

Figure 6A illustrates an anatomical view of two adjacent cross-sectional planes approximately through the primary and secondary auditory cortex. These images are assembled from the average neuronal response during the initial pre-*figure* interval (Fig. 1C). Two out of the 6 planes provided strong responses (planes 1 & 2; see **Methods**). Figure 6B displays the average responses from these two cortical slices imaged mostly during the *figure* epochs (6-12s after stimulus onset). Both 4 and 8-tone *figures* were tested, with the 8-tone evoking significantly stronger responses in both cortical planes. The evolution of the responses throughout the stimulus, including the *figure* epochs, and the subsequent *post-figure* interval is depicted in Figure 6C. All responses are referenced to the average of the first 4s interval (i.e., referenced to the 1^st^ and 2^nd^ panels from the left). The *figure* responses emerge relatively rapidly within about 1s after onset, peak within 3s, and decay in the last 2s epoch of the *figure*, a rising-falling pattern that resembles the dynamics of the single-unit responses of Figure 5. The increased 8-tone *figure* (but not 4-tone *figure*) responses persist during the *post-figure* epoch, remaining above the initial 4s of the *pre-figure* which is consistent with the single-unit findings that STRF enhancements can be estimated from the *pre*- and *post*-*figure* epochs.

Similar response patterns were measured in a second animal (ferret **Z**) whose images are shown in Figure 7. This animal was significantly older (6 years) than ferret **U** (2 years). There was a clear enhancement of responses with bigger *figures* as in the younger animal (**Figs. 6**). The notable difference between the two cases is the significantly slower dynamics of buildup and decrease in the older animal, taking about 2-3 s longer to reach its peak (Fig. 7).

**Figure 7.**
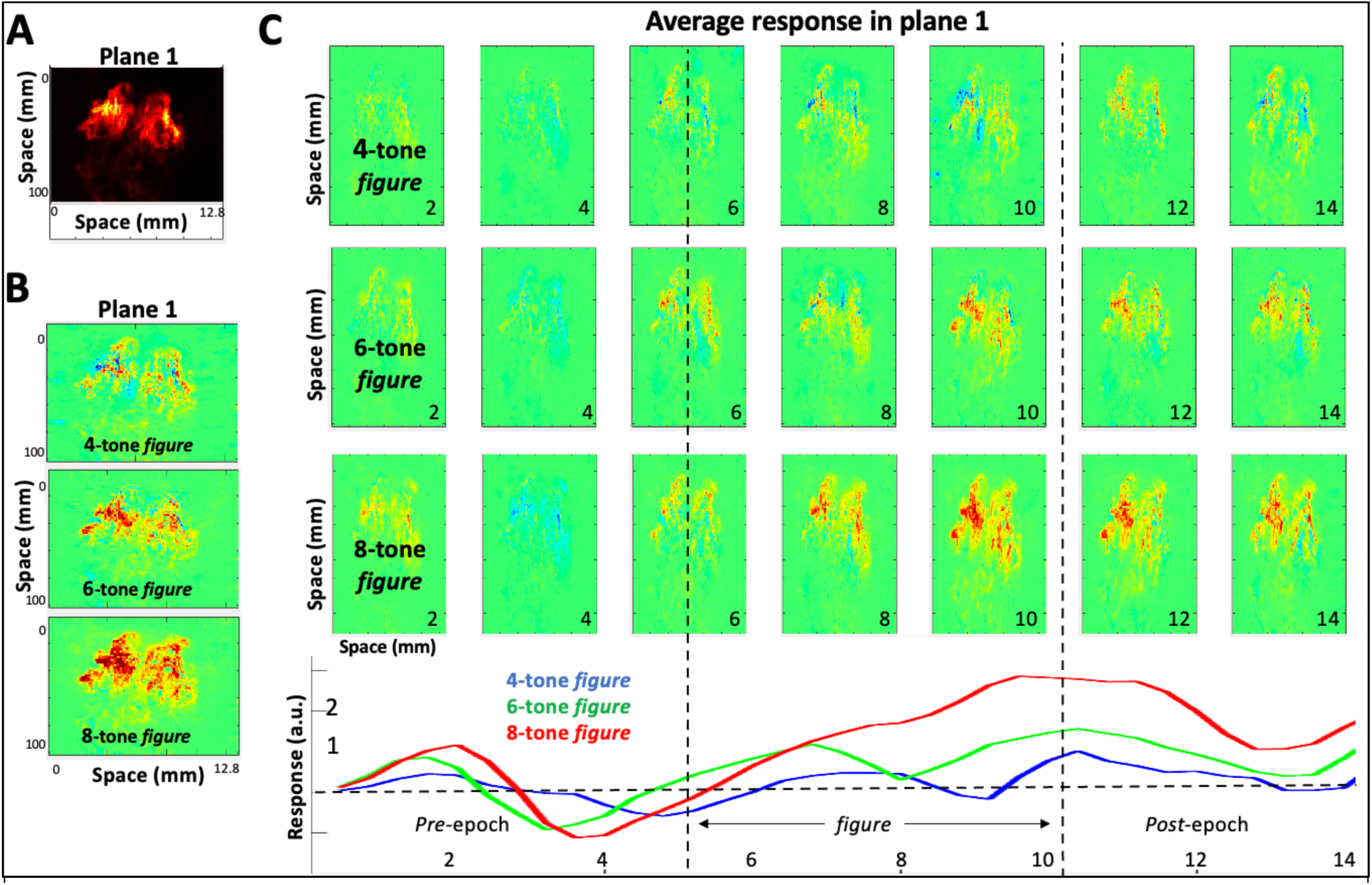
fUS imaged responses to SFG stimulus in an older ferret Z. **(A)** Anatomical vasculature of the auditory cortex in plane 1, measured by the activity induced during the early random tone-cloud. The image is the center of the primary auditory cortex, spanning 350-500 μm in thickness. **(B) Activations averaged over the entire presentations of three *figure* presentations.** Activations gradually increase with the size of the *figures*. **(C) Dynamics of the response modulations.** Sequence of 2s average activations throughout the 3 types of trials. They are strongest for the 8-tone *figures*, compared to the other two *figures*. Responses however peak at about 5 seconds from onset of the figure, significantly delayed compared to the younger animal (Fig. 6C). They decay in the latter part of the *figure* and post-*figure* epoch as in previous tests.

## DISCUSSION

This study addressed the neural mechanisms that underlie *binding,* the process that helps an animal segregate the sources in its surrounding auditory scene, by gluing together the myriad features that belong to one source (such as a voice’s pitch, timbre, and location), while simultaneously distinguishing them from those of other speakers in a mixture. It has been postulated over the last decade that binding (or equivalently the segregation of auditory scenes) relies on the temporal coherence of the sources’ signals, or the idea that temporally coherent sensory stimuli induce rapid plasticity among the synchronously activated neurons causing them to enhance their excitatory connectivity, while simultaneously inhibiting the responses of incoherently responding neurons (Fig. 1). Such synaptic modulations are presumed to occur rapidly, on the order of 100-200ms time constants [7,16] when an animal or a human switch their attentional focus from one source to another.

### Psychoacoustic and physiological studies of binding

As mentioned earlier, both experimental stimuli in Figures 1B **& 1C** have been explored in human psychoacoustic studies of stream segregation [1,9,14]. Our physiological responses are consistent with these behavioral findings in all aspects investigated. For example, the proportional increase in STRF enhancements and rapid dynamics with the size of the *figures* (**Figs. 5**) parallel well the higher detection accuracy and faster reaction times observed in humans [14] despite the absence of a behavioral task in our ferrets. The SFG stimulus mirrors many characteristics of signals in complex realistic settings, e.g., speech mixtures and orchestral musical streams. It is versatile in that it can readily be adjusted to different levels of difficulty by modifying the *figure* sizes, the jitter among its tones, or the density of the background tones. Thus, human subjects’ perception of this stimulus has been shown to reflect their performance on the Quick Speech-in-Noise tasks as well as individual differences in working memory capacity and self-reported musicianship [14].

While temporal coherence phenomena have already been demonstrated in physiological recordings in EEG/MEG/ECoG human studies [10,11,17,18,21,33], the animal experiments here afforded us a far more detailed assessment of the relationship between binding and the changes in neuronal responses and their dynamics under different attentional conditions [7]. For instance, **Experiment I** (Fig. 1B) deployed for the first-time *multiple* simultaneous streams (sequences) of different tokens, e.g., tones and noise-bursts, that could be perceptually re-organized on a trial-by-trial basis in different ways depending on how the animal selectively attends. In **Experiment II (**Fig. 1C), coherent tone-sequences (4-to 10-tone *figures*) were embedded in a complex random auditory background that modulated binding efficacy among the *figure* tones facilitating assessment of its limits and dynamics even in the absence of selective attention.

### Evidence of binding

A direct physiological test of the binding hypothesis (Fig. 1) can conceivably be done by direct observations and measurements of connectivity between cortical cells in *in-vitro* slices while undergoing synchronous *versus* asynchronous pulsatile stimulation, an experiment that has not been reported. Clearly, such experiments are difficult using an *in-vivo* preparation such as our behaving ferrets. Instead, we have measured the postulated effects of synaptic modulations among neurons driven coherently or incoherently, e.g., by measuring the dynamics and strength of their responsiveness and STRFs. Thus, in **Experiment I**, it is demonstrated that attending to the noise sequence significantly suppresses responses of neurons that are incoherent with it relative to coherent neurons which remain unaffected or become weakly enhanced (**Figs. 2B**). The role of attention in this process is critical since without favoring a neuronal response by attending to it, the two populations of coherent and incoherent responses would remain comparable and competitive and hence may not exhibit significant changes. In **Experiment II**, selective attention is absent. Instead, the balance of coherent responses (induced by the synchronized tones of the *figure*) far outweighs the incoherent responses driven by the random tones, and thus the most effective binding is that due to the *figure* coherent responses, which explains the response enhancements (Figs. 4 & 5).

### Suppression *versus* enhancement in binding

When the animals behaved in Experiment I, suppression of incoherent responses was significant in all tests, whereas enhancement relative to the passive state was often statistically weak. Nevertheless, despite the variability of the plasticity effects, the *relative* response changes were consistent with evidence for the binding hypothesis in three ways: (i) response changes in ASYN (D) versus SYN (Δ) conditions significantly satisfied the inequality **D > Δ** (Fig. 2E); (**ii**) During behavior, the binding hypothesis predicts that *contrast* between coherent and incoherent responses should increase, as found by the significant inequality **C_act_** > C_pass_ (Fig. 2F); (**iii**) ASYN and SYN response changes (**D, Δ**) in each cell were coordinated such that randomizing this association or setting **Δ** = 0 across all neurons disrupted both inequalities (**D > Δ** and **C_act_** > C_pass_ in **Figs. 2SE and 2SF**). Therefore, while the enhancements (**Δ**) in Experiment I were small (Fig.2C), they were significant enough to affect the binding as measured by its consequences.

Finally, there was a significant number of neurons that exhibited no response changes. One reason is that for binding to occur in our paradigm (Fig. 1B) neurons had to be proximate enough to interact (e.g., nearby BF’s), but far enough to avoid strong direct stimulation by both tones. The same applied to the noise stream which had to be far enough to avoid stimulating the isolated cells, but not too far to be irrelevant to the cells since binding effects are expected to weaken with distance [21]. These compromises were often difficult to balance given the independent microelectrodes. While the positioning of the tones was carefully selected relative to the BF of one of the neurons, there were undoubtedly other confounds emanating from the interaction between the stimulus tones/noise and the excitatory/inhibitory fields of the cell which may cancel out changes in the neural response.

In Experiment II, the balance of enhancements and suppression was the opposite, especially for the largest *figure* sizes (**Figs. 4**). Suppression was still evident when we focused on the plasticity in the **-ve** cells (Fig. 5B), but the dominance of such suppressive influences seen in Experiment I is absent. This is likely because of the use of sizable 6- to 10-tone *figures* that recruited and synchronized many neurons (**+ve** cells; Fig. 3C) thus inducing response enhancements. All these effects were measured in passive animals (as was the case in some of the human experiments [8–10]), but we conjecture, based on all our previous experience [7,19–23], that the binding plasticity would be significantly stronger if the SFG paradigm included a behavioral task.

### Dynamics of the binding process

Both enhancements and suppression were evident in Experiment II, where response dynamics occurred rapidly, within fractions of a second (**Figs. 5B**), suggesting that binding modulates already existing interneuron connectivity. While the onset of response modulations was rapid, it was slow to peak taking approximately 3 seconds after stimulus onset (Figs. 3B, 5), and did not persist for long as it waned although remaining elevated for seconds afterwards (**Figs. 4**, **5**). It is unclear how these details reflect a passively listening animal, as it is likely that an attentive animal may exhibit different and even faster temporal evolutions. For instance, in humans attending to and segregating speech mixtures [16], binding of different neuronal populations to change the listener’s focus and select the desired target must be significantly faster than seen here.

### Binding and aging

Since binding is essential to facilitate listening in cluttered environments, then if the process weakens with age, it becomes presumably more difficult to isolate a target voice from a mixture even if hearing remains objectively normal by most measures. The fUS measurements in the older ferret **K** (**supplementary** Figure 6S) illustrate this point in that while the *figure* enhancements appeared robust, the dynamics of the process were noticeably slower. Aged human listeners have been shown to exhibit difficulties detecting low-rate stochastic FM modulations [34] as well as poorer speech comprehension in noise, much like the SFG subjects in [14]. If confirmed in more aged animals, this stimulus may well serve as a model for the mechanisms that cause the deterioration in the binding process [24–26].

## Conclusion

Binding of temporally-coherent stimulus features is the fundamental process that ties together all human and animal psychoacoustic and physiological threads that we addressed in this study. More critically, temporal-coherence provides an overall theoretical framework to predict and interpret all the results. And, despite the absence of direct evidence to demonstrate binding as modulations of synaptic efficacy between co-activated cells, the indirect physiological consequences remain compelling as illustrated here and in previous experiments with simpler stimuli [7]. Future studies however would benefit from experiments on two ends of the spectrum. On one extreme, one may measure the direct connectivity between cell pairs or ensembles in a slice as they are stimulated coherently or incoherently. On the other extreme are experiments to engage animals in segregating and selecting a target voice in speech mixtures while recording the effects of binding across the cortical spectral channels of the target and distractor speakers [27]. Simulations of these processes [28] have already proven their efficacy at segregating complex sound mixtures. Thus, despite their relative conceptual simplicity, binding through temporal coherence may well underlie a substantial part of our remarkable abilities at disentangling sources in cluttered environments.

---

**Supplementary Figure 1S.**
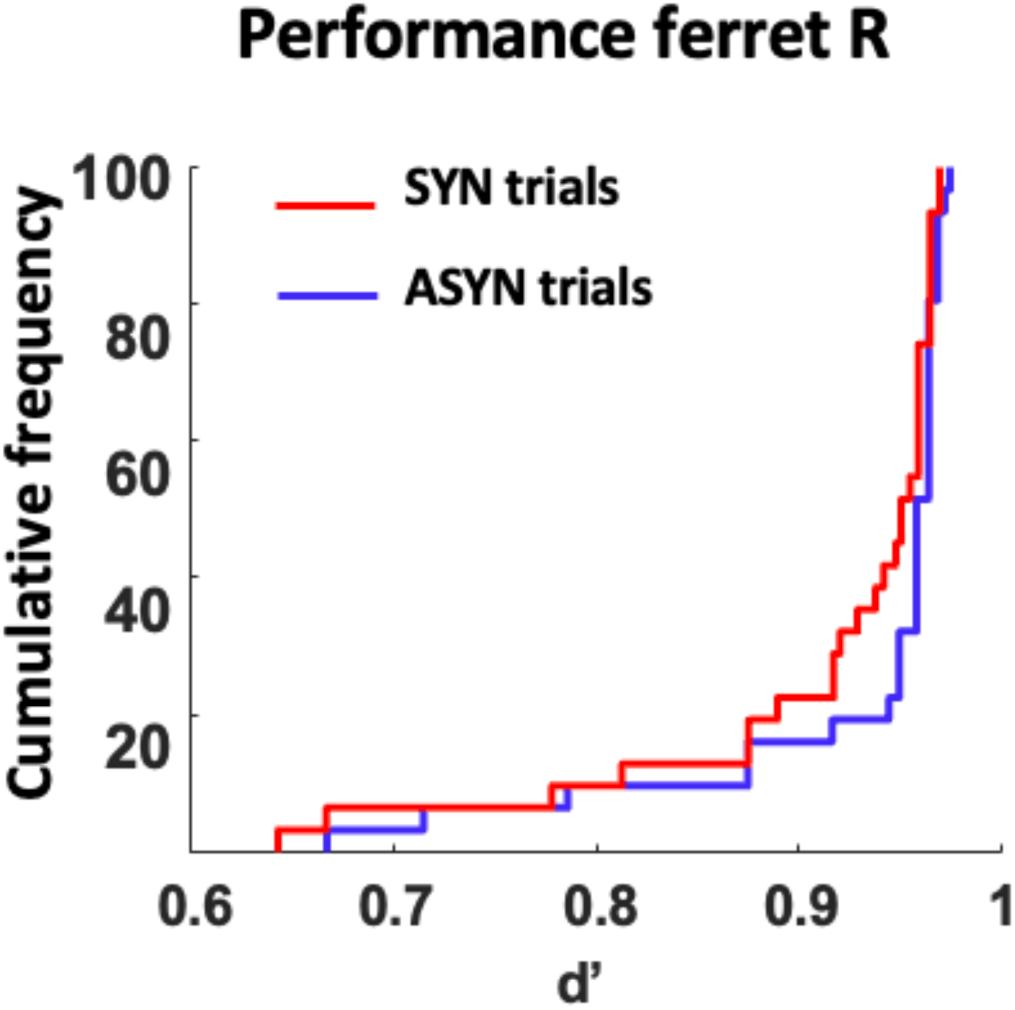
Behavior of the ferrets in Experiment I. Ferrets detected the noise deviant equally well in both SYN and ASYN trials, with no significant difference (Wilcoxon signed-rank test: p = 0.095).

**Supplementary Figure 2S.**
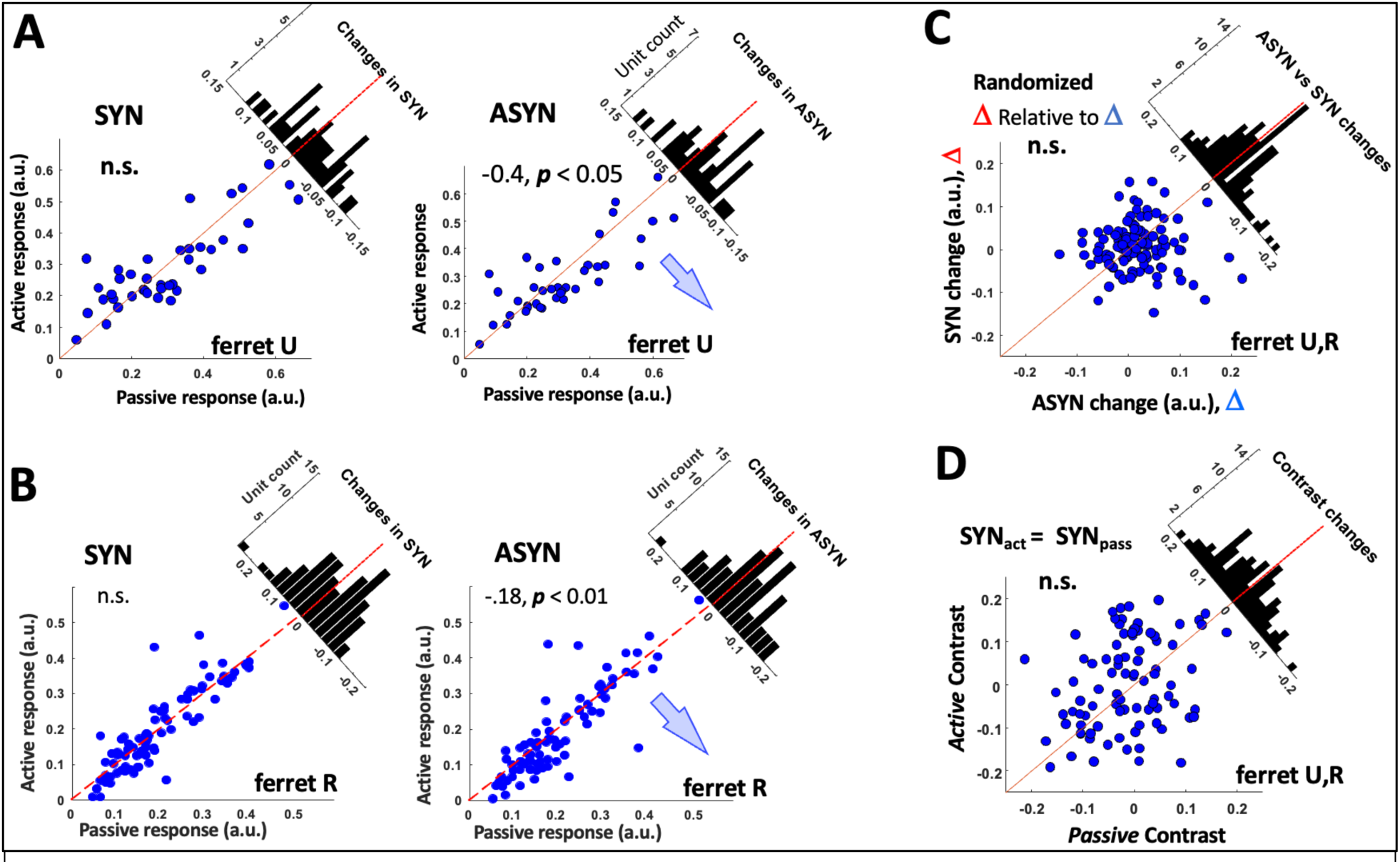
(A,B) Ferret U and Ferret R scatterplots. In each, we compare passive *versus* active for ASYM and SYM sequences. Responses are significantly suppressed in ASYM, but not significantly enhanced in SYM. (**C,D**) The SYN changes Δ, while small, do contribute to the significant biases in the scatterplots indicating binding and contrast enhancement. If the Δ contributions are randomized relative to D as explained in the text, the bias in the scatterplots of **Fig. 2D** disappears. In (**D**) **Setting** Δ **= 0** causes the *contrast enhancement* (**Fig. 2E**) to disappear, signifying the importance of Δ even if relatively small.

**Supplementary Figure 3S.**
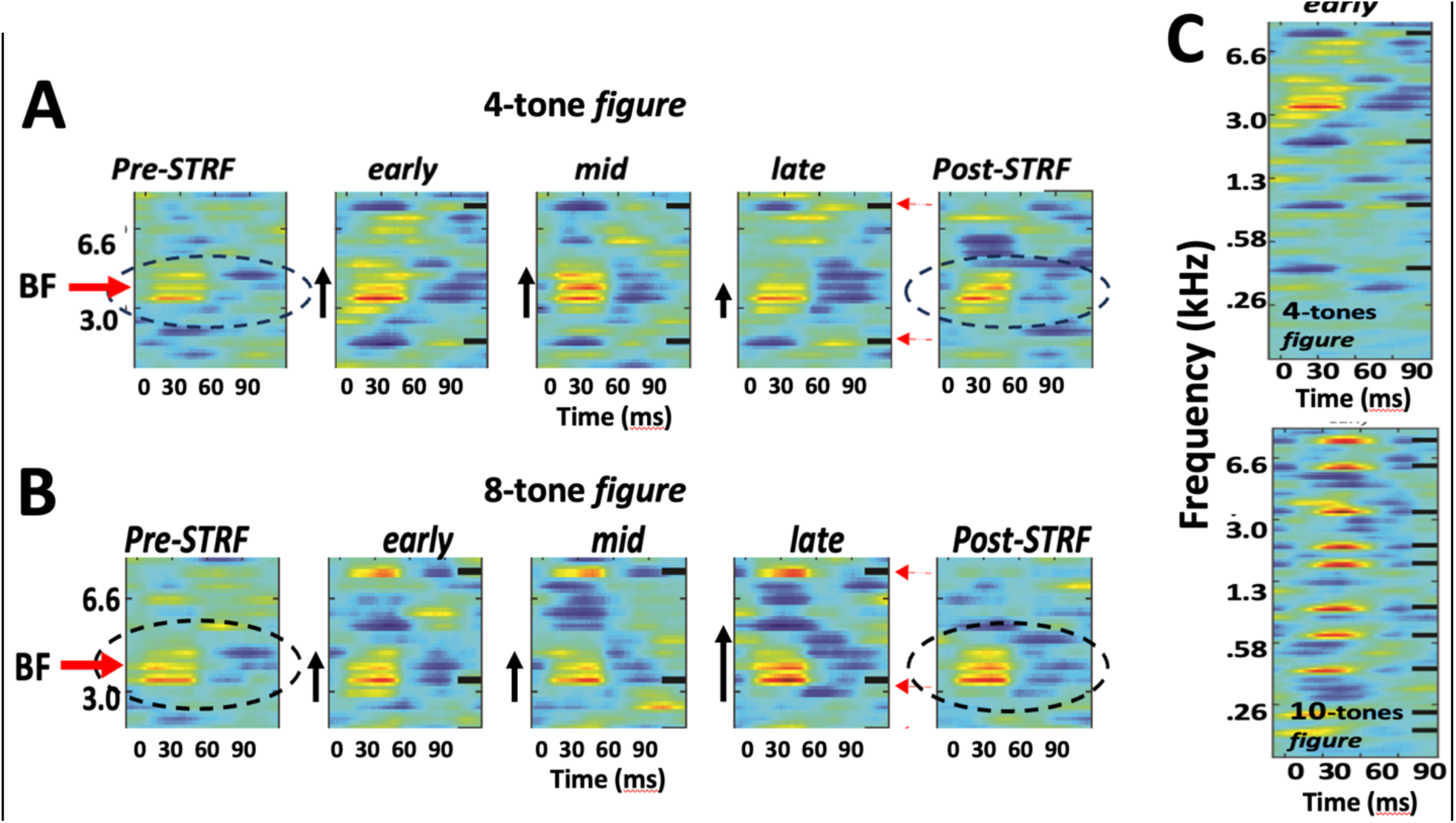
Enlarged STRFs from Figure 3. All details are identical. **(A) Partial view of the STRF series** for 4-tone *figure.* The enlargement clearly depicts the inhibitory regions aligned with the frequency of the *figure* tones (indicated by the tick marks on the right edge). Note that STRF peaks seem to decrease from earlyàlate panels. They also decrease from the *pre* à *post* panels. (B) **same as above except for the 8-*figure* tone**. Note that enhanced stripes aligned with the figure tone (the tick mark on the righthand side of each panel). This test reveals small enhancements in the STRF peaks from earlyàlate, and a small enhancement from *pre*à*post.* **(C) Full range of STRFs for 4-tone and 8-tone *figures*** depicting the inhibitory stripes (in the upper panel) and the excitatory stripes (bottom panel).

**Supplementary Figure 4S.**
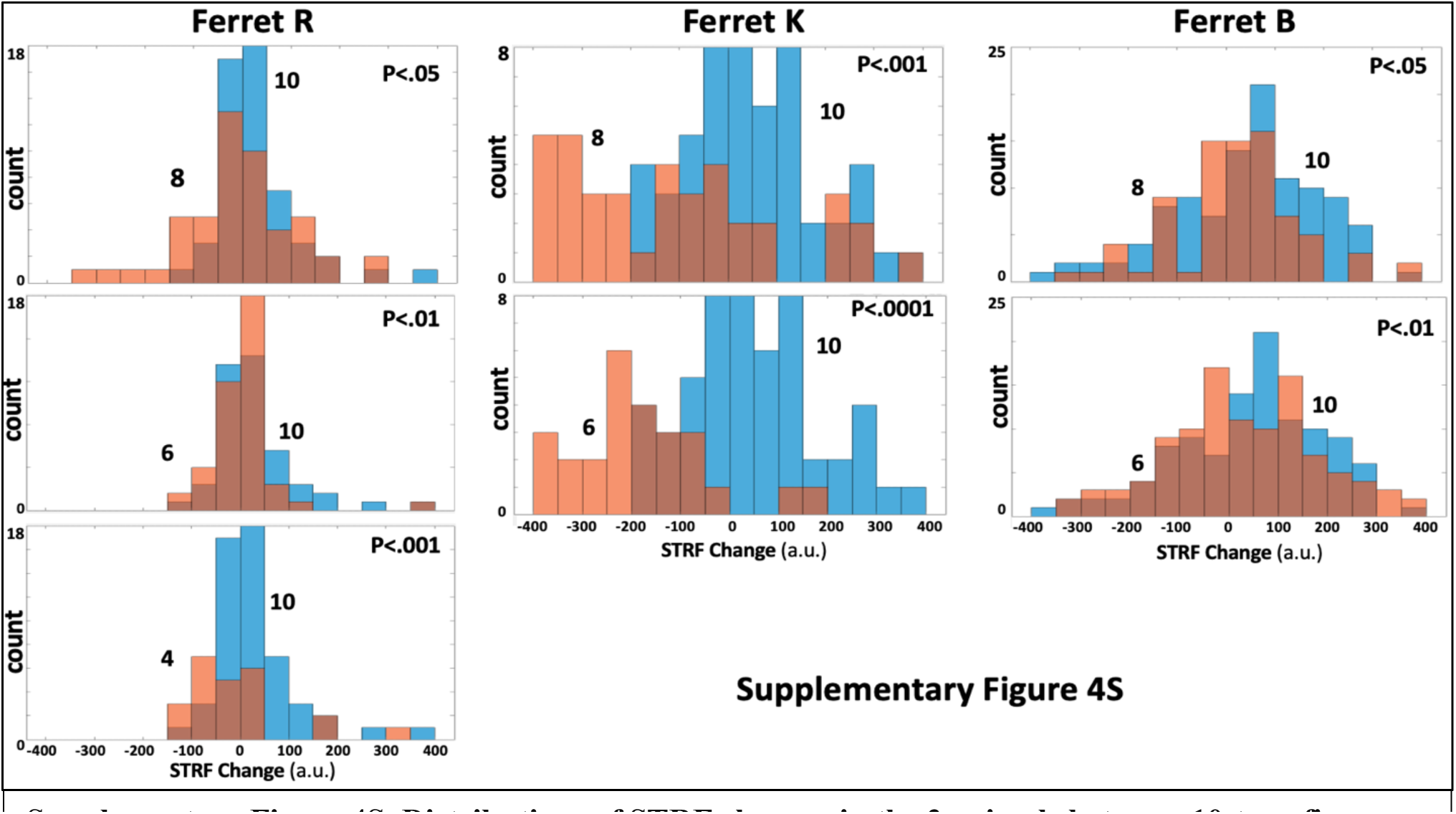
Distributions of STRF changes in the 3 animals between 10-tone figure *vs. figures* of different sizes. ***(Left column)*** Ferret R distributions for 10-tone *vs.* 8-, 6-, and 4-tone *figures. (****Middle column***) Ferret **K** distributions for 10-tone *vs.* 8-, and 6-tone *figures. (****Right column***) Ferret B distributions for 10-tone *vs.* 8-, and 6-tone *figures.* All distribution shifts to the left indicating suppression, or to the right indicating enhancements, are significant (two-sample *ttest*). Note that in all cases, the smaller *figures* exhibit more suppressed (left-shifted) STRF change relative to the 10-tone *figure* case.

## Acknowledgments

The authors would like to acknowledge funding from the National Institutes of Health (Program Grant from **NIA**, R01DC016119 **NICDC** & 1U01AG058532 **NIBI**, and a training grant **T32**).

## Author Contributions

**Kai Lu** participated in the design of Experiment I, single-unit recordings, analysis of the data, and writing of the relevant results in MS.

**Kelsey Dutta** participated in the design of Experiment II, all its single-unit recordings, and initial stages of data analyses.

**Ali Muhammed** conducted the *fUS* recording experiments.

**Mounya Elhilali** provided the conceptual guidance to the experiments, and helped with data analyses, and review of the manuscript at various stages.

**Shihab Shamma** worked on the design of the experiments, data analyses, and the writing of the manuscript.

## METHODS

### Animals

Six adult female ferrets (Mustela putorius, Marshall Farms, North Rose, NY). Two (ferrets R,U) were trained for the neurophysiological Experiment I. Three ferrets (B,U,K) others contributed to Experiment II data with the SFG stimuli in a passive state. Two more animals (ferrets U,Z) were used in the *fUS* experiments without task performance, one of which took part in Experiments I, and II.

Trained animals were placed on a water-control protocol in which they obtained water as rewards during behavioral sessions or as liquid supplements if the animals did not drink sufficiently during behavior. All animals also received ad libitum water freely over weekends, and their health was always monitored, maintaining above 80% of their ad libitum weights. Ferrets were housed in pairs or trios in facilities accredited by the Association for Assessment and Accreditation of Laboratory Animal Care (AAALAC) and were maintained on a 12-hour light-dark artificial light cycle.

All animal experimental procedures were conducted in accordance with the National Institutes of Health’s Guide for the Care and Use of Laboratory Animals and were approved by the Institutional Animal Care and Use Committee (IACUC) of the University of Maryland.

### Headpost Implant Surgeries and Neurophysiological recordings

After reaching behavioral criteria on the task, a stainless steel headpost was surgically implanted on the ferret skull under aseptic conditions while the animals were deeply anesthetized with 1% - 2% isoflurane. The headpost was secured in the skull using titanium screws and embedded in Charisma (ferrets R and U). The area around the auditory cortex was covered in a single layer of cement, whereas surrounding areas were covered with 5 – 7 mm-thick cement to protect the exposed skull.

After recovery from surgery (2 – 3 weeks), the animals were placed in a double-wall soundproof booth (IAC) and were habituated to the head-fixed setup. They were re-trained in the head-constrained version of the task. Neurophysiological recordings began after the animals regained consistent criterion levels of performance (Hit Rate = 75%, Safe Rate = 50%, Discrimination Rate = 40%) in three sequential behavioral sessions. All behavioral data in this paper were obtained after implantation, including regaining performance to pre-surgical levels.

To expose a part of the auditory cortex for recording, small 1–2 mm craniotomies were made in the skull. Recordings were conducted using 4 tungsten microelectrodes (2-5 MΩ, FHC) simultaneously advanced through the craniotomy and controlled by independently movable drives (Electrode Positioning System, Alpha-Omega) with 1 micrometer precision until well isolated spiking activity was observed. Raw neural activity traces were amplified, filtered, and digitally acquired by a data acquisition system (AlphaLab, Alpha-Omega). Single units were isolated offline using customized spike-sorting software based on PCA, K-means clustering, and subsequent template matching. Only units with greater than 70% isolation and a typical refractory period were conserved as single units.

### Localization of recording sites/ Identification of recording sites by the tonotopic

Recording locations in the primary auditory cortex were characterized by their locations along the dorso-ventral and rostro-caudal axes. Artificial marks were created by drilling a depression in the headcap on either side of the craniotomy as reference landmarks for localizing recording positions. Furthermore, the neuron’s BF was measured at each electrode penetration. The BFs obtained from all penetrations were then aligned to form a tonotopic map for all animals.

### Computing the best frequency

The best frequency (BF) of each neuron was measured by analyzing their responses to tone pips with varying frequency and intensity (see Experimental Procedures and Stimuli). A two-dimensional frequency x intensity response matrix was then created by taking the mean evoked response to tones at each frequency and intensity level. The response matrix was baseline-corrected by subtracting the mean and dividing by the standard deviation of the baseline activity from 100 ms before tone onset. The longest iso-response contour line in the normalized matrix was defined as the neuron’s tuning curve, and the frequency corresponding to the lowest intensity on the tuning curve was the neuron’s BF. A penetration site’s BF was computed by taking the median value of all isolated single units in that site.

### PSTH calculations

Peri-stimulus time histogram responses (PSTHs) were obtained by binning the neural responses into 10-ms time bins and averaging these windowed spike data across trials and stimulus presentations per cell, then further averaging across cell. Where shown, shaded error bars represented the standard error of the mean (s.e.m.) on the cell level. Unless otherwise specified, responses during engagement included only hit trials, i.e., trials where the animal successfully refrained from licking upon detecting the target change in noise level in Experiment I. PSTH responses used in further analysis are without baseline correction or normalization unless specified as “referenced to spontaneous activity,” in which case the average of neural responses during a 250-ms pre-stimulus time window was subtracted from the PSTH for each cell before averaging over population. In most plots, we have opted to show averages of PSTH of a neuronal population (unless specified otherwise). We shall refer to the response units as “**a**rbitrary **u**nits” (**a.u**.) as they become far-removed from the spike/sec rates measured with single cells.

### Experiment I

Two young female ferrets (**R,U**) were trained on an appetitive, selective attention streaming task to detect an auditory intensity cue in a sound sequence. Animals are placed on water restriction and during task performance, must withhold licking from a waterspout until the target sound cue is detected. The sound stimulus consists of alternating pure tones of 80ms duration with a cosine ramp of 5ms and 80ms gap of silence between tones. A narrowband noise (width 1/2 octave) burst plays synchronously (coherent) with one of the pure tone sequences. All reference tones and noise are at a constant intensity until the target cue occurs. After 4-30 alternating burst epochs, the target intensity cue occurs, and the animal may now lick to obtain a water reward. The target is a single noise burst played 10-20 dB louder than during the reference portion. Once they performed consistently above chance level at the final version of the task design, where consistency was defined as Accuracy (Hit Rate) = 75%, Consistent Licking pre-stimuli (Safe Rate) = 50%, and Discrimination Rate (Hit rate controlled for inconsistent licking) = 40%, we considered the animal ready for implantation. Finally, animals generally performed this task identically in the two conditions (SYN and ASYN) as exemplified by the matched performance of ferret **R** during the two sets of trials (Wilcoxon signed-rank test: p = 0.095; Supplementary Figure 1S).

### Procedures and Stimuli

Each recording began with 250-ms random tone pips of varying frequency (125 – 32000 Hz, 4 tones/octave) and intensity (0 to -50 dB range, 10 dB increment) to determine the best frequency (BF) and latency of each individual recording site. Two pure tone frequencies A and B were chosen based on the tuning properties of the neuronal units measured at a given recording depth. Effort was made to choose frequencies near one or more units’ BFs that would produce distinct responses in all units. Animals performed the task in 2 conditions: with tone A synchronous with the noise stream (*A-SYN*); and with tone B synchronous with the noise stream (*B-SYN*), or equivalently with tone A asynchronous with the noise (A-ASYN). The number of reference noise bursts *prior* to the target numbered between 4 and 20, with the exact count varied by task and pseudo-randomly trial to trial. Initially, animals performed all trials of each A-SYN and B-SYN conditions in two consecutive blocks, but later experiments randomly interspersed the two conditions. Animals had to respond within 1 second after the target onset to obtain a reward. Because the response window is longer than the target sound, 5 additional references are played after the target. Post-target reference sounds ensure that the animal is rewarded for detecting the target sound, and not simply the silence signaling the end of the trial.

Within the pre-trial reference set, tones were divided into 5-burst bins for lick count computations. Animals were allowed to lick during up to 50% of bins without triggering a terminated trial. Binning in this way prevents the animal from using a sampling strategy, that is licking once every few references to trigger a reward without listening and detecting the target. Animals were trained until they reached at least 50% discrimination rate and 50% difference in lick rate at the short trial target time. That is, if the reference counts for a trial block were 10 and 20 references, during the 20-reference trials the animal licked at the 10-reference point less than 50% of the time. This metric is to ensure that the animal is not using a timing cue and simply waiting a fixed period of seconds before licking.

Tone frequencies were chosen for each recording based on the response properties of the neurons isolated. Ideally, tones are chosen so that one but not both tones fall near the BF of each neuron. The narrowband noise is placed with center frequency at least one octave above the higher pure tone. In many experiments, noise-only trials are also played in the passive and active conditions. This paradigm represents what we hope to be a streaming task because the animal is trained to attend only to the noise, while both pure tones are distractors. However, we hypothesized that the coherent tone would be perceptually bound with the attended noise and therefore neurons responding to this tone would have enhanced responses. Analogously, neurons tuned to the asynchronous or incoherent tone were hypothesized to be suppressed. Both animals were trained until reaching threshold performance, with false-alarm trials resulting in a terminated trial and time-out session. The two animals achieved average hit rates of 93.30% ± 9.9% and 90.01% ± 11.87% with false alarm rates of 52% and 41%, respectively.

Single-unit neuronal responses were sorted and binned into 10ms windows and averaged over each 320 ms epoch of the stimulus (or two 80ms tones, each followed by 80ms gap) thus defining the PSTHs used in **Figure 2A, 2B**. Responses were analyzed from the middle third of reference epochs because we assumed that it may require a few repeats for stream formation to occur. Synchronous (SYN) and alternating (ASYN) conditions for each neuron were assigned by comparing average spike rates during passive presentation of the two alternating tone stimuli. Tone A response values in the passive state were calculated by summing tone A-noise (A-SYN, first half) with tone A alone (B-SYN, second half) responses. The same procedure was followed for tone B response values. Responses were then organized into two PSTHs labeled SYN and ASYN (**Fig. 2A**). They were assigned as SYN or ASYN *relative* to each cell based on which of these response values was greater. For example, if a neuron responds more during (A-noise + A-alone) than (B-noise + B-alone), i.e., A is the *preferred*-tone (**P** in **Fig. 2A**), then we consider A-SYN to be the SYN stimulus for that cell and B-SYN to be the asymmetric (A-ASYN) stimulus. Only neurons with a minimum 10% difference in spike rate were considered. This selection also ensured that neurons driven only by the noise stream and not tone A or B were removed. Many neurons showed changes in responses during the 4 experimental conditions (SYN_pass_, SYN_act_, ASYN_pass_ ASYN_act_). Finally, responses as expected were globally suppressed during task engagement (see text for details). However, they exhibited more suppression when driven in the ASYN_act_ condition.

### Computing the bias size around the midline

In all scatterplots of the data from this Experiment we computed a measure of the overall bias of the points around the midline. The measure captures the *effect-size* of the changes in the population. It is computed by first measuring for each point (cell or test) the signed-distance to the midline (positive/negative if above/below the midline, respectively) and then computing the mean of these signed-distances, *normalized* by the mean of the unsigned-distances. The *effect-size* therefore varies between +1 to -1 if all cells’ values fall above or below the midline.

To determine if a scatterplot bias is significant, we computed a one-sample t-test of the distribution of all distances of scatterplot points around a midline (as described above) to determine if its mean shifted significantly above or below the midline, as reflected by the indicated on each panel. If the mean shift is insignificant (*p >.05*) then both *effect-size* and its *p-values* are eliminated from the plots and the null-hypothesis (no change occurred) is validated.

### Experiment II

Three ferrets were used in the single-unit recordings of this experiment: ferret B (172), U (53), K (52) isolated single-units. Analyzed data were combined from all animals in most plots, although in some cases one animal’s data were used because of the need for consistency of stimulus parameters across all the tests. Supplemental data provide results from each animal separately. Since each cell underwent many tests with different *figure* sizes and frequency ranges, we often provided in the text and figures the more relevant total number of tests used in the analyses rather than the number of cells.

### Procedures and Stimuli

*Random tones:* A cloud of random tones (or background) consisted of a group of fixed-length (50 ms) tone pips with 5 ms cosine ramp, which occurred at random onset times within a set of discrete frequency channels. Such a cloud is generated by first selecting an octave range and a fixed number of tones logarithmically spaced per octave. For animals **B** and **U** we used 6 octaves with 6 semitones per octave, centered at 1600 Hz. For ferret **K** the same parameters were used in addition to two spectrally compressed versions, with 3 octave ranges and 12 semitones per octave, starting at the lowest frequencies of 500 and 1500 Hz. Each frequency channel had a mean tone rate of 4 Hz, i.e., on average 4 pips/s in each channel. Onset times were randomly and uniformly generated for a 2s window within a minimum of 50ms spacing between consecutive onsets, and then are jittered +/-25ms to avoid randomly generated coherence. To obtain a 5s background stimulus, we generated 6 seconds and kept the first 5, eliminating any pips that overlapped the borders.

*Perceptual Figure:* For the “*figure*” portion of the stimulus, an initial set of random onset times is generated at an average rate of 4Hz as described above. To control the precise timing of the first *figure* onset and last offset, we required that one onset occurs at t = 0 and one at t = 4.95s. Tone pips occurred in each of the *figure* frequency channels at these onset times. Then, the remaining channels were filled-in randomly as described above. Importantly, the background onsets did not overlap with the *figure* to keep the final stimulus envelope relatively flat.

*Figure* frequency channels were chosen by randomly generating a subset of 10 of the tone background channels. For the 8-tone *figure*, a subset of 8 of these 10 channels is randomly chosen, etc. for the other tone number conditions. For ease of interpretation, this procedure was performed one time, and then the same *figure* tones were used for the remainder of experiments. In most recording sessions, a set of background stimuli was pre-generated used as frozen random.

Each trial consisted of 5s of background, followed by 5s of background with an embedded figure, then 5s background. We hypothesized that the temporal coherence of the *figure* tones would cause them to be bound into a stream, and that this effect size would systematically vary with the size of the *figure*, or the number of tones which make up the *figure*. Typically, multiple *figure* sizes were generated and tested with at least 50 repetitions per *figure* size.

### Data analyses

The background of tones is an auditory stimulus akin to coarse noise, which is commonly used to assess neuronal receptive fields. We first validated that our version of this stimulus can recapitulate response fields of A1 generated by our standard tuning battery, specifically TORC responses [13]. Using lag values of -10 to 120ms, the cross-covariance of the response and stimulus (auditory spectrogram) was divided by the auto-covariance of the stimulus. Ridge regression was performed with 30 log-spaced values of *λ* and the results averaged. This procedure was performed for each trial and then averaged, using stimuli and responses to 1s windows immediately before the figure onset (*pre*), immediately after the first onset (*early*), halfway through figure presentation (*mid*), the last second of figure presentation (*late*), and immediately following the last figure onset (*post*). The background responses can reconstruct STRFs generated from TORC responses, including the classic checkerboard patterns. Previous studies have indicated that tuning maps generated with background stimuli can vary systematically with tone density, with more spectro-temporally dense clouds producing sharper tuning [29]. We chose our density parameters based on intuition of human perceptual limits, but it is possible that a denser background may produce better tuning and different effects. It is important to note also that TORCs contain a continuous range of frequencies spanning a 5-octave range, while the SFG contains only a discrete set of 37 frequencies spanning typically 6 octaves. Thus, excitatory or inhibitory regions that span multiple channels and appear smooth are in fact interpolated from a few frequency samples by the spectral blurring inherent in the auditory spectrogram.

Neurons demonstrated typical responses to SFG stimuli. Units that were tuned to frequencies far from any of the *figure* tones had relatively stable responses throughout the stimulus. Some units showed strongly suppressed responses to the *figure* tones while maintaining excitatory responses to their best frequency (BF). Still others were tuned at a frequency equal to a *figure* tone and hence showed strong excitation in response to *figure* tones. The reasons these STRF features emerge is because the *figure* tones are correlated during the *figure* presentation, and hence it is not possible to disambiguate responses driven by any one of the tones. Therefore, we examined the response field during the post-STRF period to identify any tuning modulation driven by the *figure* presentation (see text for details). Previous studies have shown that attention-dependent plasticity can persist for minutes or up to hours [19], so the one-second window immediately following the *figure* should retain some of the modulatory effects.

To test for STRF significance, we generated a null distribution of voxel values using the same procedure as described above for STRFs, but mismatching responses and stimuli. We then down-selected the 128 frequency bins in the auditory spectrogram by removing all channels which did not correspond to the pure tone frequencies in the stimulus. A Wilcoxon paired t-test was used to select voxels from the data with significant distributions (*p <* 0.05). Summing over the 37 (frequency bins) by 130 (lags) response map yielded mean correlation values for each of the 5 epochs described above (see text for details). Subtracting the *pre-figure* from the *post-figure* yields a summary value for the size and direction of change in response - stimulus correlation (see text). We also averaged over the response from 20 to 80ms lag to select the strongest response area corresponding to the BF of the isolated cell.

On a population level, we found that the degree of change between *post*-*figure* and *pre*-*figure* increased systematically with *figure* size. This is depicted in the distributions of Figure 4D in the text, which are fitted by the best-fit normal distributions for the two extreme *figures* (4- and 10-tones). For 4-tone *figure*, which is perceptually difficult for humans, the change curves centered with a low peak. For 10-tone *figures*, which produce a strong perceptual pop-out [9,11,14], the curve is right-shifted, with a peak in the positive region.

### Functional Ultrasound Imaging

Functional Ultrasound imaging (*fUS*) is a recording technology based on blood flow imaging [30]. It is a variation of Doppler-based ultrasonic imaging that measures the ultrasonic energy back-scattered from red blood cells (proportional to blood volume) in each voxel of the image. The technique is widely used in medical imaging - but major innovations have turned it into a powerful tool for neuroscience such as an ultrafast imaging scanner able to acquire thousands of images per second and filtering to remove global coherent signals [31,32]. These developments have significantly boosted signal-to-noise ratio (SNR) and increased spatial resolution to ∼100μm, allowing us to measure subtle changes in blood ow due to local neuronal activity.

We implanted two ferrets (**U**, age < 3 yrs**; Z**, age > 5 yrs) for the *fUS* recordings. Ferrets were first surgically implanted with a stainless steel headpost that was attached to the sagittal interparietal suture and secured in the skull with titanium screws and cement then areas surrounding the auditory cortices were covered with radiopaque bone cement (PALACOS), leaving 1–2 cm^2^ cavities for easy access to the auditory cortex in both hemispheres. During surgery ketamine (35 mg kg^−1^ intramuscularly) and dexmedetomidine (0.03 mg kg^−1^ subcutaneously) are used to anesthetize ferrets and 1−2% isoflurane to maintain anesthesia also atropine sulfate (0.05 mg kg^−1^ subcutaneously) was used to control salivation and to increase heart and respiratory rates. Electrocardiogram, pulse, and blood oxygenation were monitored, and rectal temperature was maintained at ∼38 °C. During surgery the skull was surgically exposed and the areas of auditory cortex were determined and marked in both side of skull. Following surgery, antibiotics (cefazolin, 25 mg kg^−1^subcutaneously) and analgesics (dexamethasone 2 mg kg^−1^ subcutaneously and flunixin meglumine 0.3 mg kg^−1^ subcutaneously) were administered. ferrets were allowed to recover for 2 weeks before being habituated to a head restraint in a customized horizontal cylindrical holder for a period of 2 weeks. Then, the next surgery for the fUS window was performed using a surgical micro drill to remove cement and skull bone, yielding relatively large craniotomies over the auditory cortex ∼ 15×10 mm window over the brain. After clean-up and antibiotic application, the hole was sealed with an ultrasound-transparent TPX cover, embedded in an implant of radiopaque bone cement (PALACOS). Sterile polysiloxane impression material (EXAMIX NDS; GC America, Inc.) was placed in the wells between recording sessions, which allowed the craniotomies to be kept well-protected from the environment. (EXAMIX NDS; GC America, Inc.) and were cleaned and treated with topical antiseptic drugs (povidone-iodine). The skin surrounding the implant was cleaned three times per week with warm saline and treated with povidone-iodine and sulfadiazine cream ointment.

*<u>Ultrasound Imaging</u>:* All acoustic stimuli were presented at 65–70 dB SPL. Sounds were digitally generated at 40 kHz with custom-made MATLAB functions and A/D hardware (PCI-6052E; National Instruments) and presented with a free-field speaker positioned 30 cm in front of the animal’s head. During imaging sessions, the ultrasonic probe was placed in contact with the implant cement and acoustic coupling was assured via degassed ultrasound gel. Experiments were conducted in a double-walled sound attenuation chamber. We used a custom miniaturized probe (15 MHz central frequency, 128 elements) inserted in a four degree-of-freedom motorized setup. The probe was driven using a stereotaxic manipulator with a custom made holder to control its position.

*<u>Imaging:</u>* Functional Ultrasound (FUS) imaging has rapid acquisition of 300 frames at a 500 Hz frame rate (lasting 600ms, that is one to two ferret cardiac cycles) that are filtered to remove tissue motion from the signal using a spatiotemporal clutter filter, each frame being a compound frame acquired via 11 tilted plane wave emissions (−10° to 10° with 2° steps) fired at a pulse-repetition frequency (PRF) of 5500 Hz. And the Image reconstruction is performed using GPU-parallelized delay-and-sum beamforming. A CBV image is averaged every second to capture the dynamics of the cerebral blood physiological response. High sampling rate is a key asset to cancel any respiratory or tissue pulsatile motion artifacts in the final averaged images.

Power doppler is then computed for each voxel (100 x 100 x ∼400 µm, over the 300 time points which is proportional to blood volume. A certain band of Doppler frequencies can be chosen before computation of the power using a bandpass filter enabling the selection of a particular range of axial blood flow speeds such as a slow blood flow of capillaries and arterioles vs fast blood flow of and big vessels. We focused on small vessels with axial velocity lower than 3.1mm.s^−1^. Power UfD signal was normalized towards the baseline to monitor changes in Cerebral Blood Volume (%CBV).

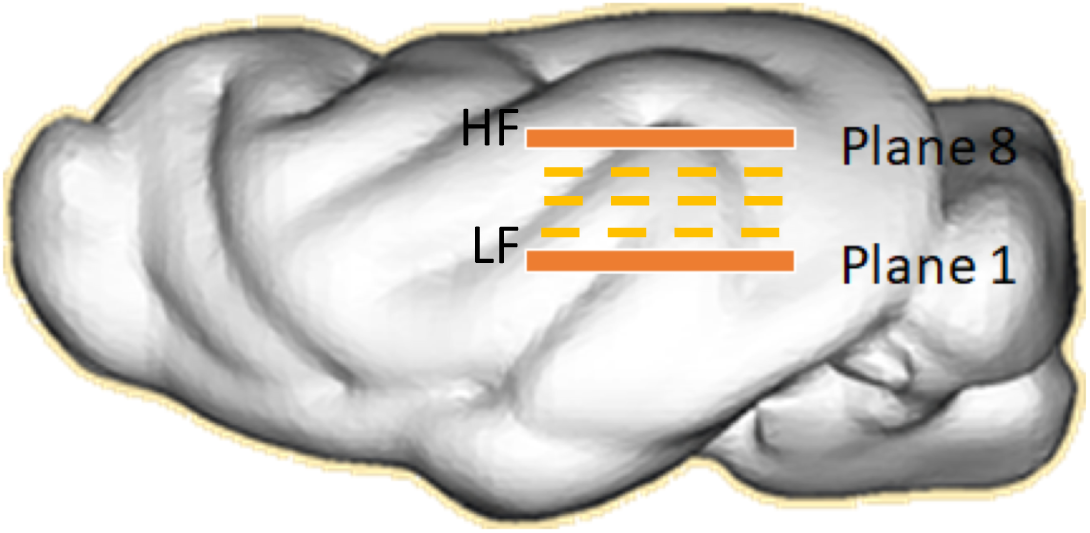

The imaging progressed by first positioning the ultrasound transducer probe at plane 1, we then ran the SFG stimulus as described above for a total of 50 repetitions. In each presentation, the stimulus consisted of a pre-*figure* (5s), *figure* period (5s), and a post-*figure* (5s), ending with silence for 5s. The probe was then moved 400um to Plane 2 and the same procedure was repeated. The process was repeated up to 6 or 8 planes as necessary.

The localization of the fUS images on the auditory cortex is based on the anatomical vasculature of the auditory cortex (e.g., Figs 6A and Fig. 7A) which provide familiar landmarks based on our previous publications [15,30,32}.

